# EXOSC10 sculpts the transcriptome during the growth-to-maturation transition in mouse oocytes

**DOI:** 10.1101/663377

**Authors:** Di Wu, Jurrien Dean

## Abstract

Growing mammalian oocytes accumulate substantial amounts of RNA, most of which is degraded during subsequent meiotic maturation. The growth-to-maturation transition begins with germinal vesicle or nuclear envelope breakdown (GVBD) and is critical for oocyte quality and early development. The molecular machinery responsible for the oocyte transcriptome transition remains unclear. Here, we report that an exosome-associated RNase, EXOSC10, sculpts the transcriptome to facilitate the growth-to-maturation transition of mouse oocytes. We establish an oocyte-specific conditional knockout of *Exosc10* in mice using CRISPR/Cas9 which results in female subfertility due to delayed GVBD. By performing multiple single oocyte RNA-seq, we document dysregulation of several types of RNA, and the mRNAs that encode proteins important for endomembrane trafficking and meiotic cell cycle. As expected, EXOSC10-depleted oocytes have impaired endomembrane components including endosomes, lysosomes, endoplasmic reticulum and Golgi. In addition, CDK1 fails to activate, possibly due to persistent WEE1 activity, which blocks lamina phosphorylation and disassembly. Moreover, we identified rRNA processing defects that cause higher percentage of developmentally incompetent oocytes after EXOSC10 depletion. Collectively, we propose that EXOSC10 promotes normal growth-to-maturation transition in mouse oocytes by sculpting the transcriptome to degrade RNAs encoding growth-phase factors and, thus, support the maturation phase of oogenesis.

## INTRODUCTION

Improper oocyte maturation directly causes ovulatory disorders and leads to female infertility (1,2). Maturing oocytes dramatically alter their transcriptome in preparation of fertilization and early embryonic development. Due to the absence of transcription, active RNA degradation plays a vital role in transcriptome remodeling. During this transition, many transcripts are deadenylated which uncouples the translation machinery from the poly(A)-binding complex and exposes both 5’ and 3’ ends to RNA degradation (3,4). Certain dormant RNAs, including *Mos, Plat* and *Cnot7*, have short or no poly(A) tails and are stable during oocyte growth (5,6). After resumption of meiosis, these RNAs are rapidly polyadenylated to translate meiosis-required proteins. This is apparently triggered by exposure of their untranslated regions (7) which are captured by specific RNA binding proteins (5). A growing body of work documents that coordinated sequestration, deadenylation, translation and degradation collectively regulate RNA metabolism during oocyte maturation. A dysregulated transcriptome can impair oogenesis and post-fertilization embryogenesis (4,8-12).

For historic reasons, the oocyte nucleus is referred to as the germinal vesicle (GV). After the growth phase, GV-intact oocytes remain quiescent in pre-ovulatory follicles until induced to mature by luteinizing hormone. Critical events in the transition from growth to maturation include chromatin condensation, termination of transcription and germinal vesicle breakdown (GVBD). The first two are directly coupled and associated with increased repressive histone modifications (13,14). The different timing and order of chromatin changes result in two distinct DNA configuration, namely NSN (non-surrounded nucleolus) and SN (surrounded nucleolus) GV-intact oocytes, having less or more developmental potential, respectively. The nuclear configuration also exhibits distinct status of nucleolar rRNA processing (15). The third event, GVBD, requires precise coordination of the meiotic cell cycle and membrane trafficking. Meiotic cell cycle control is determined by cyclin B/CDK1 activity through complex signaling pathways including cAMP-PKA and PKB/Akt (16). High levels of cAMP cause CDK1 inhibitory phosphorylation that favors GV arrest. Conversely, decreased cAMP activates CDK1 to form the functional cyclin B/CDK1 complex that induces GVBD. The membrane trafficking machinery involves multiple endomembrane components. For example, an exchange apparatus of ER-NE (endoplasmic reticulum - nuclear envelope) appears to facilitate NE formation post mitosis by reorganizing membrane structures around chromatin (17). The COPI-coated vesicles that normally traffic from the Golgi to the ER also promote GVBD upon recruitment by nucleoporin Nup153 (18). However, the molecular basis coordinating the transcriptome with GVBD events is not known.

The RNA exosome is a highly conserved complex that degrades or processes cellular RNAs from the 3’ end. RNA exosome-related genetic mutations have been identified in a wide range of diseases, including diarrhea of infancy, neurodegenerative disorders and multiple myeloma (19-21). The association of the core complex with DIS3, DIS3L or EXOSC10 RNases provide the required enzymatic activity. EXOSC10 is a nuclear RNase, the absence of which causes RNA processing defects in yeast (22) and increased sensitivity to DNA damage in fly and human cells (23,24). EXOSC10 has been documented to promote mRNA turnover (25), 3’ pre-rRNA processing (26) and long noncoding/enhancer RNA degradation (27). The depletion of EXOSC10 in human cell lines can stabilize short-poly(A) RNAs and increase the length of their poly(A) tails (28) which confirms that EXOSC10 normally functions to degrade RNA post deadenylation. It also has been reported to control the onset of spermatogenesis in male germ cells (29). However, whether EXOSC10 is essential for oogenesis has remained unclear. Published data sets (4,30) document that *Exosc10* transcripts are highly abundant in mouse oocytes and early embryos, raising the possibility of its participation in maternal RNA metabolism.

In the current study to evaluate EXOSC10 function in oogenesis, we established oocyte specific conditional *Exosc10* knockout (cKO) mice. cKO females had decreased fertility associated with defects in GVBD and impaired oocyte maturation. Using single oocyte RNA-seq with ERCC spike-in normalization, we identified dysregulated genes involved in endomembrane trafficking and CDK1 phosphorylation. Other than protein coding genes, there are also defects in snoRNA and rRNA in cKO oocytes, suggesting additional mechanisms of EXOSC10 function consistent with data from cell lines. These observations suggest a major role of RNA degradation in sculpting the transcriptome of maturing, transcriptionally quiescent oocytes prior to activation of the embryonic genome.

## MATERIALS AND METHODS

### Mice

Mice were maintained in compliance with the guidelines of the Animal Care and Use Committee of the National Institutes of Health under a Division of Intramural Research, NIDDK-approved animal study protocol.

### Generation of *Exosc10* floxed allele by CRISPR/Cas9

Two guide RNAs (gRNA, 50 ng/μl), two homology-directed repair (HDR, 100 ng/μl) templates and *Cas9* cRNA (100 ng/μl) were microinjected into 1-cell mouse embryos. The two gRNA sequences were: 5’tcagtggagacctgcgatct3’ (left loxP) and 5’gaaattctgatgtctagcgg3’ (right loxP). Each gRNA was transcribed *in vitro* from double stranded (ds) DNA templates with a MEGAshortscript™ T7 Transcription Kit (Thermo Fisher Scientific, AM1354) and purified by a MEGAclear Transcription Clean-Up Kit (Thermo Fisher Scientific, AM1908). The dsDNA templates for sgRNAs were initially synthesized as single stranded (ss) DNA by Integrated DNA Technologies (IDT) and amplified by primers 5’GATCCCTAATACGACTCACTATAG3’ and 5’ AAAAAAAGCACCGACTCGGTGCCAC3’ into ds DNA: 5’gatccctaatacgactcactataggtcagtggagacctgcgatctgttttagagctagaaatagcaagttaaaataaggctagtccgttatcaacttgaaaaagtggcaccgagtcggtgcttttttt3’ and 5’gatccctaatacgactcactatagggaaattctgatgtctagcgggttttagagctagaaatagcaagttaaaataaggctagtccgttatcaacttgaaaaagtggcaccgagtcggtgcttttttt3’.

The two HDR templates for each edited locus were synthesized as ssDNA by IDT: 5’gagagagcacgtatggctcttgcagaggactggtactctaccccagcacccatgttaggtgggtcacaactgcttgtaactccagctccaagaGCGGCCGCATAACTTCGTATAATGTATGCTATACGAAGTTATtcgcaggtctccactgacactggcactcaggagcacgt3’ (left loxP) and 5’actcacactgtagaccagtctggcctcaaactcacaaagatccacctgcctctgcctcctaagtgctggggttaaatgggtactctaccaccgGAATTCATAACTTCGTATAGCATACATTATACGAAGTTATctagacatcagaatttctaaatataaaaaggagaatg3’ (right loxP).

After linearization with Pmel, plasmid #42251 (Addgene) was used as a template to transcribe *Cas9* cRNA *in vitro* by a mMESSAGE mMACHINE™ T7 ULTRA Transcription Kit (Thermo Fisher Scientific, AM1345). The synthesized cRNA was purified by a MEGAclear Transcription Clean-Up Kit (Thermo Fisher Scientific, AM1908).

For mouse embryo microinjections, hormonally stimulated B6D2_F1_ female mice were mated to B6D2_F1_ males. 1-cell zygotes were flushed from oviducts into M2 medium (CytoSpring, #M2114) and microinjected with the mixed components for gene-editing with CRISPR/Cas9. The injected embryos were cultured in KSOM (CytoSpring, #K0113) at 37 °C with 5% CO_2_ for 24 hr. 2-cell embryos were transferred into the oviducts of 0.5-day post coitus pseudopregnant ICR females.

To obtain *Exosc10* oocyte-specific conditional KO mice (cKO), *Exosc10* floxed mice were crossed to *Zp3-cre* mice (31). The genotyping primers for the *Exosc10* floxed and deletion alleles were as follows:

left loxP: 5’atgagtcgggtaatgcagtac and 5’tgtgtgaggatgttgtgagc3’;

right loxP: 5’ccgactctgacattgagtgg3’ and 5’gcctctttcccacagttccag3’;

Deletion allele: 5’atgagtcgggtaatgcagtac and 5’gcctctttcccacagttccag3;

Cre: 5’gcggtctggcagtaaaaactatc3’ and 5’gtgaaacagcattgctgtcactt3’.

### Oocyte collection and culture

GV oocytes collection: ovaries were dissected from female mice (6-10 weeks old) into M2 medium plus milrinone (2.5 μM). The ovaries were pierced mechanically by 30-gauge-needles to release oocytes and only fully-grown oocytes (GV oocytes) detaching easily from granulosa cells were collected for further experiments. For *ex vivo* oocyte maturation, GV oocytes were washed (20 times) with M2 medium without milrinone and cultured in M2 at 37 °C with 5% CO_2_. The GVBD/GV ratio was determined at GV+3h (GV3h) and meiosis II progression was evaluated at 14 hr.

Growing oocytes collection: ovaries were dissected from female mice (7-10 days old) into M2 medium plus milrinone. The ovaries were punctured by 30-gauge-needles to release follicles and growing oocytes. The nude growing oocytes were collected for further experiments.

### cRNA *in vitro* transcription and microinjection

The *Exosc10* coding sequence was inserted into plasmid #44118 (Addgene) to form an in-frame fusion with mVenus. Simultaneously a T7 promoter (TAATACGACTCACTATAGGG) was inserted into the 5’ end of the *Exosc10* coding sequence. The plasmid was linearized by XbaI, purified and *in vitro* transcribed with a mMESSAGE mMACHINE™ T7 ULTRA Transcription Kit (Thermo Fisher Scientific, AM1345). The cRNAs were purified with a MEGAclear Transcription Clean-Up Kit (Thermo Fisher Scientific, AM1908) and diluted to a proper concentration (500 ng/μl, unless otherwise stated) for microinjection.

The microinjection station consisted of a Zeiss inverted microscope, a pair of Eppendorf Transferman NK2 micro-manipulators and an Eppendorf Femtojet 4i Injector. The microinjection needles were made from borosilicate glass (with outer diameter of 1.0 mm) using a Sutter Flaming/Brown Micropipette Puller. The injection solution (cRNA or CRISPR/Cas9 injection mix) was loaded into the injection needle from the back by Eppendorf Microloader and the holding pipet was purchased from Eppendorf (VacuTip).

During microinjections, the oocytes were kept in 150 μl M2 (2.5 μM milrinone) medium on a glass slide. The injection time (ti) was set at 0.1 s, the compensation pressure (Pc) was set as 15, and the injection pressure (Pi) was varied (100-500) depending on different needle shapes and flow to ensure that the injected volume was ∼1-2 pl.

### Immunofluorescence and confocal microscopy

*Ex vivo* cultured oocytes and embryos were fixed in 2% paraformaldehyde (Thermo Fisher Scientific, #50-980-492), diluted in PBS containing 0.1% Triton X-100 at 37 °C for 30 min. After fixation, oocytes were washed with PBVT (PBS, 3 mg/ml polyvinylpyrrolidone-40 and 0.1% Tween-20) and permeabilized with 0.5% Triton X-100 in PBS for 30 min at room temperature. Oocytes were blocked with 5% normal goat serum in PBVT for 1 hr at room temperature and then incubated with primary antibody overnight at 4 °C. On the second day, the oocytes were washed (4X, 15 min) with PBVT and incubated with fluorescence conjugated secondary antibody overnight at 4 °C. On the third day, after washing with PBVT (4X, 15 min), oocytes were stained with DAPI and mounted with PBS for confocal microscopy (LSM 780, Carl Zeiss).

### Source and dilution of antibodies, staining reagents and live staining dyes

Anti-GM130 (1:200, BD Transduction Laboratories 610823); LysoTracker™ Green DND-26 (1:1000, ThermoFisher L7526); LysoTracker™ Blue DND-22 (1:1000, ThermoFisher L7525); ER tracker red (1:1000, Invitrogen E34250); anti-lamin B1 (B-10) (1:200, Santa Cruz sc-374015); anti α–tubulin (1:200, Sigma T5168); anti-lamin A/C (4C11) (1:200, CST 4777T); anti-phospholamin A/C (Ser22) (1:200, CST 2026); anti-RAB5 [EPR21801] (1:500, Abcam ab218624); anti-phosphoCDK1 (Thr14, Tyr15) (17H29L7) (1:200, ThermoFisher 701808); anti-cAMP (1:200, RD # MAB2146); anti-pericentrin (1:2000, Abcam #ab4448); anti γ–tubulin (1:500, Abcam ab11316); anti-rRNA antibody (Y10b), (1:100, Novus NB100-662SS); goat anti-rabbit IgG (H+L) cross-adsorbed secondary antibody, Alexa Fluor 546 (1:500, Invitrogen A-11010); goat anti-mouse IgG (H+L) cross-adsorbed secondary antibody, Alexa Fluor 647 (1:500, Invitrogen A-21235). DAPI (Sigma D9542-1MG); goat serum (Sigma G9023-10ML); Tween-20 (Sigma P1379-25ML).

### Ovary histology

Ovaries were dissected from female mice into PBS buffer. After removing surrounding lipid and tissue, the ovaries were fixed in newly prepared 2.5% glutaraldehyde and 2.5% paraformaldehyde in 0.083 M sodium cacodylate buffer (pH 7.2) for 3-5 hr at 4 °C. Ovaries were washed with 0.1 M sodium cacodylate buffer (pH 7.2) and kept at 4 °C overnight. Finally, the ovaries were transferred into 70% ethanol and stored for more than 1 day at 4 °C. Sectioning (2 μm, every 20^th^ section), periodic acid–Schiff (PAS) staining and mounting were performed by American HistoLab, Gaithersburg, MD. The imaging of ovary histology was performed on a Nikon ECLIPSE Ti microscope using a 20x objective.

### RNA FISH with oligo(dT) probes

FISH assays were performed in U-bottom 96-well plates. Oocytes were fixed with 2% paraformaldehyde (Thermo Fisher Scientific, #50-980-492), diluted in PBS containing 0.1% Triton X-100 at 37 °C for 30 min. Oocytes were dehydrated stepwise at room temperature using 25%, 50%, 75%, 100% (methanol: PBS volume) and washed (3X) with 100% methanol. After - 20 °C treatment >1 hr, the oocytes were rehydrated into PBT (PBS-0.1% Tween-20) through 75%, 50%, 25%, 0% (methanol: PBT volume) steps. Oocytes then were washed (3X) with PBT, treated with 0.5% SDS for 30 min at room temperature and re-washed (3X) with PBT. Oocytes were washed (2X) with freshly prepared FISH wash buffer (10% formamide, 2x SSC, 0.1% Tween-20 in nuclease-free water), transferred into hybridization buffer (10% formamide, 2x SSC, 10% dextran sulfate in nuclease-free water) and incubated at 37 °C for 2 hr. The oocytes then were incubated with probe-containing hybridization buffer for 16 hr at 37 °C. On the second day, oocytes were incubated sequentially with hybridization buffer (30 min, 37 °C), FISH wash buffer (1X, 5 min; 4X, 20 min at 37 °C), and FISH wash buffer (1X, 20 min) at room temperature. Oocytes were further washed (2X) with PBT at room temperature, mounted onto ProLong™ Diamond Antifade Mountant (Thermo Fisher Scientific, P36970) and imaged by confocal microscopy (LSM 780, Carl Zeiss).

### Translation activity detected by Click-iT HPG assay

Translation activity was detected by a Click-iT® HPG Alexa Fluor® 594 Protein Synthesis Assay Kit (Thermo Fisher Scientific, C10429). Briefly, oocytes were cultured in M2 medium containing HPG (50 μM) for 30 min, washed twice with PBS, fixed with 2% paraformaldehyde, washed twice with 3% BSA-PBS and detected with a Click-iT HPG kit per the manufacture’s protocol. The oocytes then were imaged by confocal microscopy (LSM 780, Carl Zeiss) and processed with ImageJ (Fiji) software.

### Single oocyte poly(A)-based RNA-seq libraries

Single oocyte RNA-seq libraries were prepared according to a published single-cell RNA-seq pipeline with minor modifications (32). Briefly, oocytes at desired stages were collected individually into 2.5 μl RLT Plus (Qiagen) and stored at −80 °C until all samples were acquired. In the beginning of RNA purification, an ERCC RNA spike-in mix (Thermo Fisher Scientific, 4456740) was diluted 10^5^ fold, and 1 μl of the diluted ERCC mix was added to each sample. Poly(A) RNA was isolated by oligo (dT) beads, reverse transcribed, amplified and purified. Preliminary sequencing was performed with different amplification cycles (10, 12, 14, 16, 18) and different portions of oocytes as initial material (1/8, 1/4, 1/2, 1) to test the linear range of amplification. 14-cycles was chosen as the best condition based on the cDNA acquired and the regression analysis of ERCC. The purified cDNAs were analyzed by Bioanalyzer 2100 (Agilent) to confirm successful amplification and quality. Qualified cDNAs were used to construct sequencing libraries by Nextera DNA Sample Preparation Kits (Illumina). The resultant 71 libraries were evaluated by a Bioanalyzer 2100 and pooled into 6 groups for purification. The sequencing was performed by the NIDDK Genomic Core Facility using the HiSeq 2500 Sequencing System (Illumina).

### Single oocyte RiboMinus RNA-seq libraries

Single oocyte RiboMinus RNA-seq libraries were prepared with the Ovation® SoLo RNA-Seq Library Preparation Kit (#0501-32) after minor modifications. Briefly, 1 μl of 10^5^-fold diluted ERCC RNA spike-in mix (Thermo Fisher Scientific, 4456740) was added to each oocyte lysis sample. For first strand cDNA synthesis, 1 μl (50 μM) of random hexamer primer (Thermo Scientific 3005) was added to each sample instead of the provided First Strand Primer Mix (blue: A1 ver 17). In the library amplification I step, 16 cycles were used based on qPCR optimization. The purified cDNA libraries were analyzed by a Bioanalyzer 2100 (Agilent) to confirm successful amplification and quality. The sequencing was performed by the NIDDK Genomic Core Facility using the HiSeq 2500 Sequencing System.

### Single oocyte rRNA sequencing libraries

rRNA libraries were prepared along with RiboMinus RNA-seq libraries described above without ribosome depletion (AnyDeplete). All libraries were pooled in one lane and sequenced by the NIDDK Genomic Core Facility using the HiSeq 2500 Sequencing System.

### RNA-seq analysis based on ERCC RNA spike-in mix

Raw sequence reads were trimmed with Cutadapt 2.5 to remove adapters while performing light quality trimming with parameters “-m 10 -q 20, 20”. Sequencing library quality was assessed with FastQC v0.11.8 with default parameters. Trimmed reads were mapped to the Mus musculus mm10 reference genome plus ERCC.fasta using STAR v2.7.2a. Multi-mapping reads were filtered using samtools 1.9. Uniquely aligned reads were then mapped to gene features using HTseq v0.9.1 as an unstranded library with default parameters. A gene/ERCC count was considered valid when present in at least 5 reads in at least 2 libraries. Differential expression between groups of samples was tested using R version 3.5.1 (2018-07-02) with DESeq2 v1.24.0. The oocyte libraries were normalized by ERCC counts by defining the ERCC genes as the controlGenes when estimating the sizeFactors in DESeq2. The functional annotation of the genes of interest was performed using the DAVID website. Note that all RNA-seq differential analyses used ERCC normalization (except Figure 2I which used median ratio normalization) which corrected for potential deviations due to differences in library size. For intronic and intergenic read analyses, gtf files were produced by BED tools and reads were directly counted against the special gtf files.

**Figure 1.**
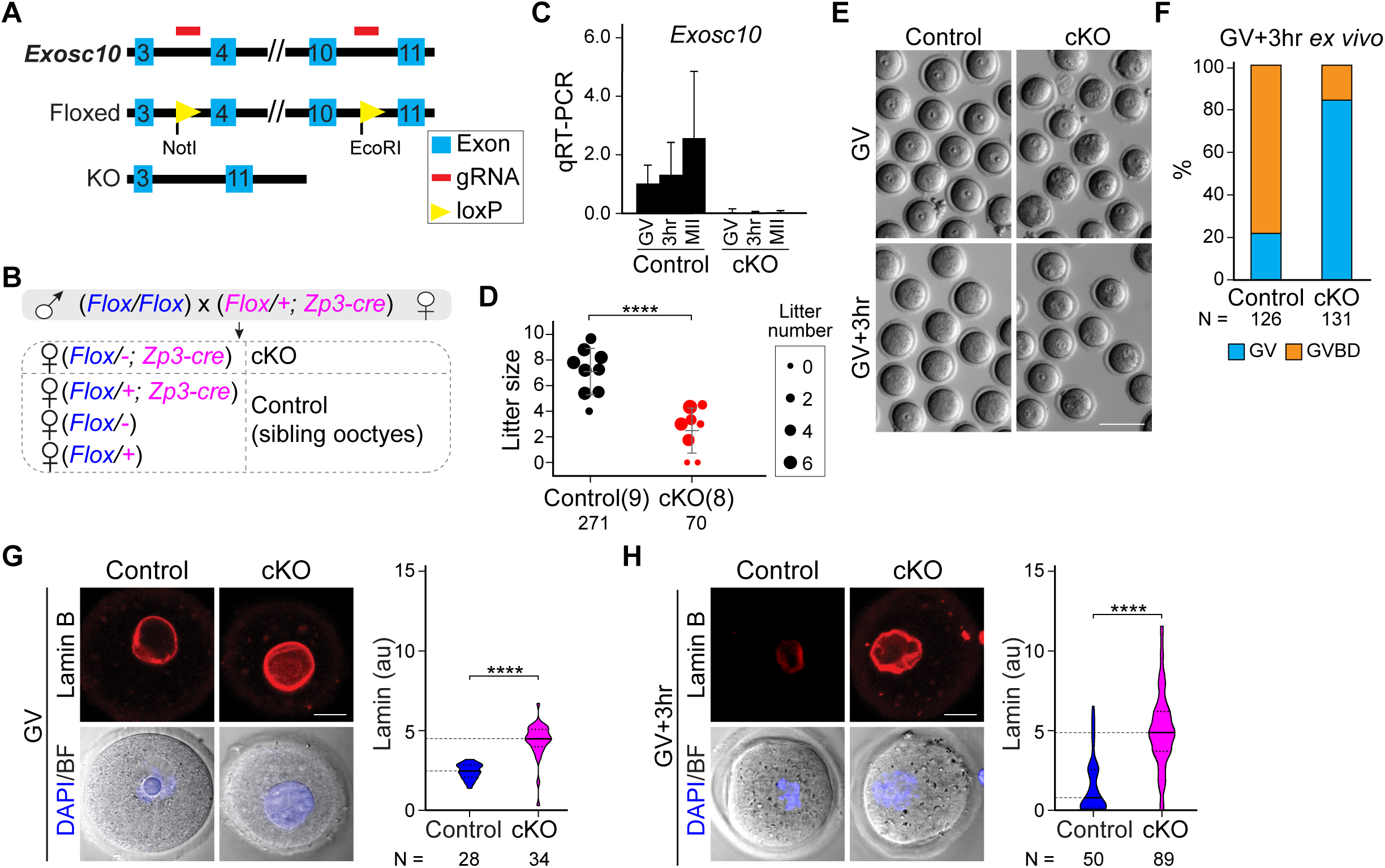
Oocyte-specific knockout of *Exosc10* causes female subfertility by impairing germinal vesicle breakdown during oocyte maturation. (**A**) Schematic of strategy to generate an *Exosc10* floxed allele using CRISPR/Cas9. Two loxP sites were inserted to bracket exons 4-10. (**B**) Mating strategy to obtain oocyte-specific conditional knockouts of *Exosc10* (cKO). Siblings with other genotypes were used as controls. The paternal allele is labeled in blue and the maternal allele is labeled in magenta in the offspring. Note that the floxed maternal *Exosc10* allele will become a deletion allele (-) during oocyte growth. (**C**) qRT-PCR of *Exosc10* in single oocytes obtained from controls and cKO mice. (**D**) Dot plot of individual litter sizes over 6-months of harem breeding of controls and cKO females with wildtype males. The sizes of the dots are normalized by the average litter number per female. The number in parenthesis is the number of females having the indicated genotypes. The number of pups born is indicated below each group. The horizontal lines represent the mean and standard deviation. (**E**) Bright-field images cKO and controls oocytes cultured *ex vivo* for 0 (GV) or 3 hr (GV3h). (**F**) Percentage of GVBD oocytes in **E**. Number of oocytes indicated below each group. (**G-H**) Confocal fluorescence and brightfield/DAPI images of oocytes after lamin B immunostaining at GV (G) and GV3h (H) stages. Lamin B and DAPI are maximum intensity projections and bright-field images are single optical sections containing the nucleus. Quantification of lamin B fluorescence is on the right. The horizontal lines inside the violins represent the median and the quartiles. The number of oocytes from at least 3 females are indicated below each group. **** *P* <0.0001 in **D, G, H**, two-tailed Student’s t-test. Scale bars: 100 μm in **E**; 20 μm in **G, H**.

**Figure 2.**
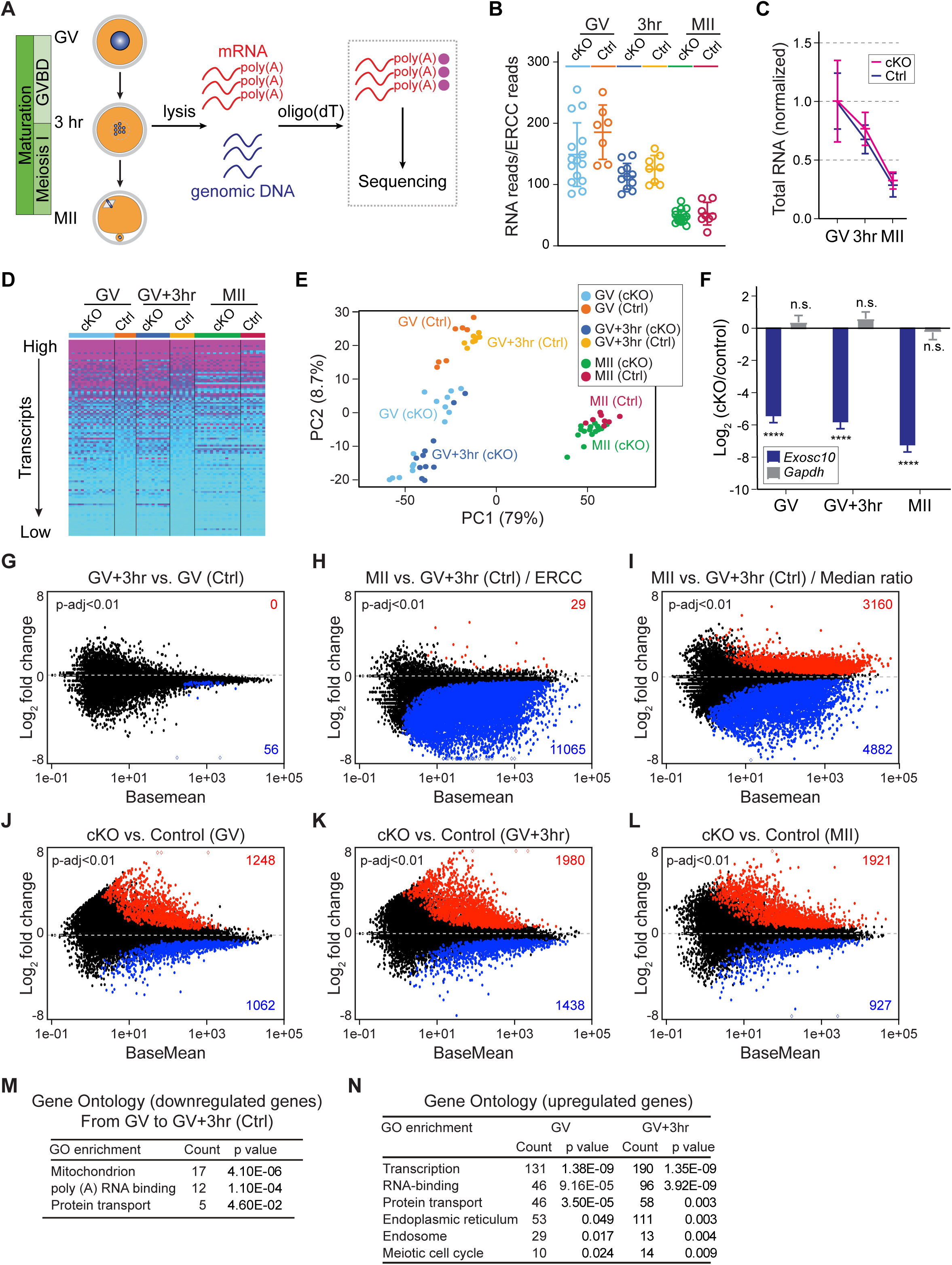
*Exosc10*^*cKO*^ oocytes exhibit dysregulated transcriptomes during oocyte maturation. **(A)** Schematic illustrating the pipeline of single oocyte RNA-seq. After individual oocyte lysis, oligo-dT beads captured poly(A) RNAs for library construction and sequencing. Genotypes were determined from genomic DNA. (**B**) Total RNA levels were normalized for each library by an ERCC RNA spike-in mix. (**C**) Further normalization of total RNA level in **B** by the mean value of the GV stage within each genotype. (**D**) Heatmap of all libraries, each row represents one gene and each column represents one library. Genes are ranked from highest to lowest expression level, and every 100^th^ gene from the top half were selected to represent the transcriptome. The transcription level was color-coded from high to low. (**E**) Principal component analysis (PCA) of the 64 libraries. Each dot represents one library, color coded by genotype and stage. (**F**) Log_2_ fold change of *Exosc10* and *Gapdh* in cKO vs. control oocytes. The bars and lines are log_2_ fold change and standard error of the mean from DESeq2 analyses. **** *P*-adjust <0.0001, n.s. no significance, which are the *P*-adjust values in DESeq2 analysis. (**G-H**) MA-plots of transcript changes from GV to GV3h, and from GV3h to MII stage in control oocytes. The more and less abundant transcripts in each comparison are labeled by red and blue, respectively (both have *P*-adjust <0.01). (**I**) MA plot of transcript change from GV3h to MII stage in control oocytes when analyzed by median ratio normalization. (**J-L**) MA-plots of transcript changes in cKO vs. control oocytes at GV, GV3h and MII stages. The increased and decreased abundant transcripts in each comparison are labeled by red and blue, respectively (*P*-adjust <0.01). (**M-N**) Gene ontology of the more abundant transcripts in **G, J** and **K**.

### Usage of the published data set (GSE70116)

Using published RNA-seq data sets of growing oocytes (GO) and fully-grown oocytes (FGO) (12), we selected the median value at each stage to define if transcripts had increased or decreased abundance. Then we defined transcripts into GO-specific, FGO-specific, GO-FGO increased and GO-FGO decreased groups based on relative abundance levels between GO and FGO.

### Quantitative RT-PCR

Single oocytes were collected individually for poly(A) RNA enrichment, purification, reverse-transcription and amplification following a similar pipeline as the single oocyte RNA-seq protocol. The purified cDNA was used directly as templates and the qRT-PCR was performed by iTaq Universal SYBR Green Supermix (Bio-Rad, #1725121) and QuantStudio 6 Flex Real-Time PCR System (Thermo Fisher Scientific). The primers for *Exosc10* and *Gapdh* were:

*Exosc10* primer 1: 5’ccgactctgacattgagtgg3’ and 5’gcctctttcccacagttccag3’;

*Exosc10* primer 2: 5’ATCCCCCAGGGAAAGACTTC3’ and 5’GTCCGACTTTCCAACAGCAA3’;

G*apdh*: 5’TGCACCACCAACTGCTTAGC3’ and 5’GGCATGGACTGTGGTCATGAG3’.

### Statistical analysis

The replicates of the oocytes, ovaries or mice were combined and analyzed by paired-sample t-test, which was performed by GraphPad Prism 8 software. The fluorescence quantification was performed with ImageJ (Fiji) software.

## RESULTS

### Oocyte-specific knockout of *Exosc10* leads to female subfertility by disrupting oocyte maturation

Homozygous constitutive ablation of *Exosc10* in mice is embryonic lethal (Figure S1A, C). To study the function of EXOSC10 in oocyte development, we generated an *Exosc10* floxed allele by CRISPR/Cas9 and used *Zp3-cre* mice to specifically knock out *Exosc10* in growing oocytes. Two loxP sites surrounding exons 4-10 resulted in a deletion and a frame shift of the remaining coding sequence (Figure 1A; Figure S1B). To increase loxP recombination efficiency, we used a mating strategy to present only one floxed allele for cre to obtain the desired oocyte-conditional knockout (*Flox/-; Zp3-cre*) which was designated cKO (Figure 1B). qPCR of single oocytes confirmed the decrease of *Exosc10* transcripts in cKO mice beginning early in the growth phase and extending beyond the GV stage (Figure 1C; Figure S1B, D).

To assess female fertility after oocyte specific EXOSC10 depletion, we mated pairs of cKO and control females with wildtype males for 6 months. Combining the records of 7 harems including 8 cKO females and 9 controls, cKO females had substantial subfertility with reduced total number of pups (8.8 vs. 30.1), number of litters (2.6 vs. 4.1) and litter size (3.3 vs. 7.3) per female (Figure 1D). Intrigued by the subfertility, we investigated possible defects in oocyte growth, maturation and early embryogenesis. At 12 weeks, cKO and control females had indistinguishable ovaries in terms of weight, histology and number of antral follicles (Figure S1E-H) which suggests normal oocyte growth. Although the diameter of the cKO oocytes was modestly decreased (Figure S1I), we concluded that the subfertility of cKO females is likely due to defects after the growth phase.

Next, we examined oocyte maturation and determined that GV oocytes collected from cKO females had deficient meiotic progression after 20 hr *ex vivo* culture (Figure S1J). Correspondingly, the number of ovulated eggs recovered following gonadotropin stimulation also was significantly decreased in cKO females (Figure S1K). We narrowed the defect to GVBD (Figure 1E-F). The microtubule-organizing center proteins appeared normal in the cKO oocytes, suggesting normal initiation of spindle formation (Figure S1L). However, lamin B persisted at GV+3hr (GV3h) in cKO oocytes, and its intensity was substantially increased at both GV and GV3h stages compared to controls (Figure 1G-H). Using *in vitro* synthesized *Exosc10-mVenus* cRNA, we found that EXOSC10-mVenus had nuclear localization in both oocytes and embryos (Figure S1M), which could pass into the cytoplasm post GVBD. We also evaluated pre-implantation development of embryos derived from homozygous cKO females. In *ex vivo* culture of 1-cell embryos, many had developmental delay from embryonic day 1.5 (E1.5) to E3.5. Some of them arrested at the 2-cell stage and there was decreased progression to blastocysts at E3.5 (Figure S1N-O). In sum, oocyte-specific EXOSC10 depletion caused substantial subfertility due to defective GVBD and embryos derived from cKO females progressed abnormally during pre-implantation development.

### EXOSC10 depletion dysregulates poly(A) RNA profiles

To investigate the molecular basis of this phenotype, we sought to identify the types of RNA regulated by EXOSC10 in oocytes. EXOSC10 exhibits ribonuclease activity toward a range of poly(A) RNA molecules, including mRNA and noncoding RNA (33,34). We therefore employed RNA FISH to quantify the poly(A) RNA with an oligo(dT) probe. In wildtype oocytes, there was a modest decrease during maturation from GV to GV3h and then a sharp decrease during progression to MII (Figure S2A-B) which is consistent with the reported RNA degradation during oocyte maturation (4). Oocytes, over-expressing *Exosc10-mVenus*, had significantly decreased poly(A) compared to *mVenus* alone over-expression at both GV and GV3h stages, suggesting a role for EXOSC10 in degrading poly(A) RNA. This was substantiated by mutating the ribonuclease catalytic sites (D313N and E315Q) in EXOSC10 after which over-expression of the *dExosc10-mVenus* lost the ability to accelerate poly(A) RNA degradation (Figure S2C-D). On the other hand, the overall poly(A) RNA intensity in cKO oocytes showed no obvious change compared to controls (Figure S2E-F). These results indicate that EXOSC10 participates in poly(A) RNA metabolism during mouse oocyte maturation.

To characterize the dysregulated poly(A) RNA that accounts for the phenotype, we performed single oocyte RNA-seq at GV, GV3h and MII stages. The poly(A) RNA were isolated by oligo-dT beads (Figure 2A) and an equal amount of ERCC Spike-In Mix was added to each oocyte lysate to ensure library quality and to compare initial RNA quantities (Figure S3). In total, 71 oocytes (from 7 controls and 8 cKO mice) were sequenced, and 64 passed quality control. In normalizing oocyte total RNA with ERCC, we observed dependence on library size (Figure 2B). In addition to an overall mild delay in RNA degradation (Figure 2C), cKO oocytes exhibited distinct gene expression patterns as determined by a pan-transcriptome expression heatmap (transcripts were ranked by expression level and every 100^th^ was plotted) and principal component analysis (Figure 2D-E; Table S1). As expected, the near complete loss of *Exosc10* transcripts in cKO oocytes was validated (Figure 2F).

ERCC not only provides a stringent criterion for library quality evaluation, but also allows comparison among libraries that vary considerably in size. Within control oocytes, we identified 56 transcripts (*P*-adjust <0.01) with decreased abundance in GV and GV3h. From GV3h to MII stage, we identified 11,065 transcripts with decreased abundance. There were almost no transcripts with increased abundance which is consistent with minimal transcription during oocyte maturation (Figure 2G-H). In contrast, the default median-ratio normalization without ERCC revealed both increased and decreased abundance of transcripts in comparing GV3h and MII stages, presumably due to the influence of library size on normalization (Figure 2I). Thus, ERCC normalization was used in subsequent comparisons of transcript abundance between stages and genotypes. At each stage, we identified more transcripts with increased than with decreased abundance in cKO oocytes (Figure 2J-L; Table S1). We performed GO analysis of the transcripts that differed in abundance from GV to GV3h in control groups and the transcripts that differed in abundance in cKO oocytes. The most prominent terms were associated with transcriptional control, RNA metabolism, endomembrane transport and meiosis (Figure 2M-N). All of these biologic processes play important roles in nuclear envelope breakdown (35) which links the dysregulated transcriptome in cKO oocytes with the failure of GVBD.

### Heterogeneity of the cKO oocytes underlies subfertility

The RNA FISH and the PCA of the RNA-seq suggest high similarity between GV and GV3h stages of control oocytes and more variation in the cKO oocytes. It seemed plausible that greater dysregulation of the transcriptome would correlate with more reduced fertility of individual cKO oocytes. Thus, we performed PCA again using the 41 samples of control oocytes and cKO oocytes at both GV and GV3h stages and employed k-means clustering to define three groups (group number k=3). As expected, there was one group that included all the control oocytes which is consistent with the high similarity between GV and GV3h of control oocytes analyzed in the original PCA (Figure 2E). The two other groups were designated cKO (major) and cKO (minor) according to their greater and lesser distances from the control group on the PCA plot (Figure 3A-B). By differential analysis of cKO (major) and cKO (minor) vs. control, we found that the cKO (major) group exhibited more higher abundant transcripts than the cKO (minor) group (Figure 3C-D). Those transcripts that differ in abundance between cKO (major) and cKO (minor) groups were enriched for GO terms including transcription regulation, mitochondrion, cell cycle and endomembrane vesicles, which recapitulated the observed differences in the original cKO vs. control (Figure 3E-F; Figure 2N; Table S1). To confirm the stronger phenotype in cKO (major) compared to cKO (minor), we examined six transcripts that were markedly most abundant in the 6 GO terms enriched by cKO vs. control (Figure 2N): *Zfp668, Hormad1, Nox4, Xpot, Exoc8* and *Dio3*. The abundance of transcripts from all six genes was significantly higher in the cKO (major) compared to the cKO (minor) group (Figure 3G). Thus, greater dysregulation of the transcriptome in individual oocytes decreased their potential fertility.

**Figure 3.**
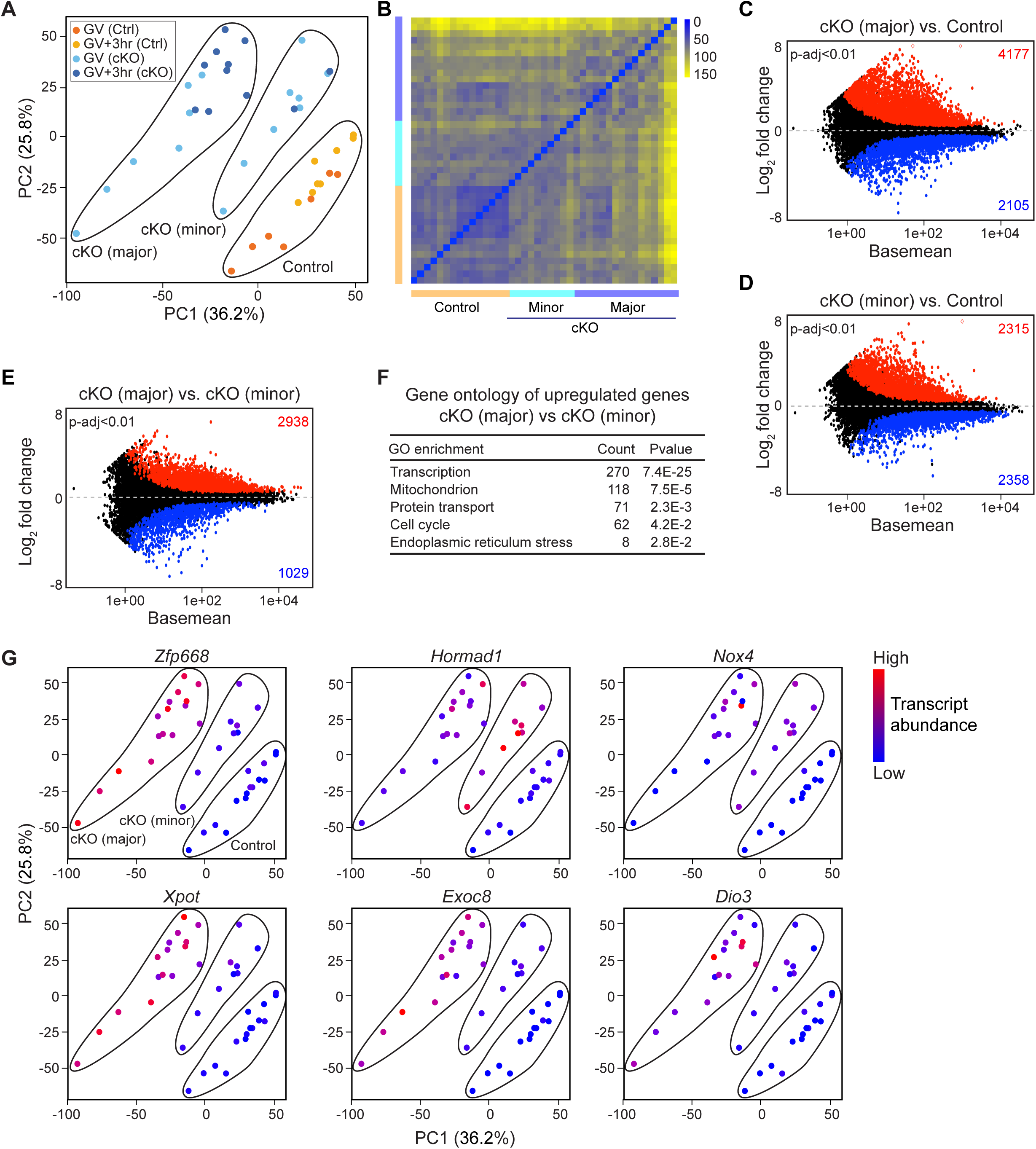
Transcriptome heterogeneity of *Exosc10*^*cKO*^ oocytes underlies the subfertility. **(A)** Principal component analysis (PCA) of 41 libraries of control oocytes and cKO oocytes at both GV and GV3h stages. Subsequently the 41 libraries were sub-clustered by k-means algorithm into 3 groups: control; cKO (major); and cKO (minor). Dots are color-coded to indicate library source. (**B**) Sample distance matrix of the 41 libraries in **A**. (**C**-**D**) MA-plots of cKO (major) vs. control and cKO (minor) vs. control libraries. (**E**) MA-plot of cKO (major) vs. cKO (minor). The increased and decreased transcripts in each comparison are labeled by red and blue, respectively (*P*-adjust <0.01). (**F**) Gene ontology of the more abundant transcripts with log_2_ fold change more than 1 in **E**. (**G**) PCA plots of **A** that is color-coded by the expression level of 6 different transcripts in individual oocytes. The control, cKO (major) and cKO (minor) groups in each plot are the same as labeled in the first plot.

### EXOSC10 depletion results in non-poly(A) RNA dysregulation

During oocyte maturation, polyadenylation occurs when mRNA is being translated, and deadenylation occurs when mRNA is being degraded or stored (6). To rule out the potential interference of the dynamic poly(A) status to poly(A)-based RNA-seq and to capture non-poly(A) transcripts, we performed single oocyte RiboMinus RNA-seq. Due to the high similarity between GV and GV3h stages, we didn’t include the GV3h in the RiboMinus RNA-seq. The ERCC normalization of GV oocytes (8 from 2 control females, 7 from 2 cKO females) and MII eggs (4 from 2 control females, 3 from 2 cKO females) indicates a decrease of transcriptome size during oocyte maturation, which is impaired in cKO oocytes (Figure 4A-B; Figure S4). Similarly, the heterogeneity of the cKO oocytes at the GV stage could be visualized through the PCA clustering (Figure 4C). When comparing MII to GV in the control group, we identified 12,220 less abundant transcripts (*P*-adjust <0.01), which is similar to comparing MII and GV3h in the poly(A)-based RNA-seq results (Figure 4D; 2H). In cKO oocytes, there were 2,563 more and 124 less abundant transcripts at the GV stage, and 10,081 more and 89 less abundant transcripts at the MII stage (*P*-adjust <0.01). The changed transcripts could enrich similar functional terms as those from poly(A)-based RNA-seq, including transcription, RNA-binding, protein transport, ER, mitochondrion, etc. (Figure 4G; Figure 2N; Table S2). The more abundant transcripts in cKO samples were scattered across the entire coding region indicating the absence of partial degradation (Figure S5A).

**Figure 4.**
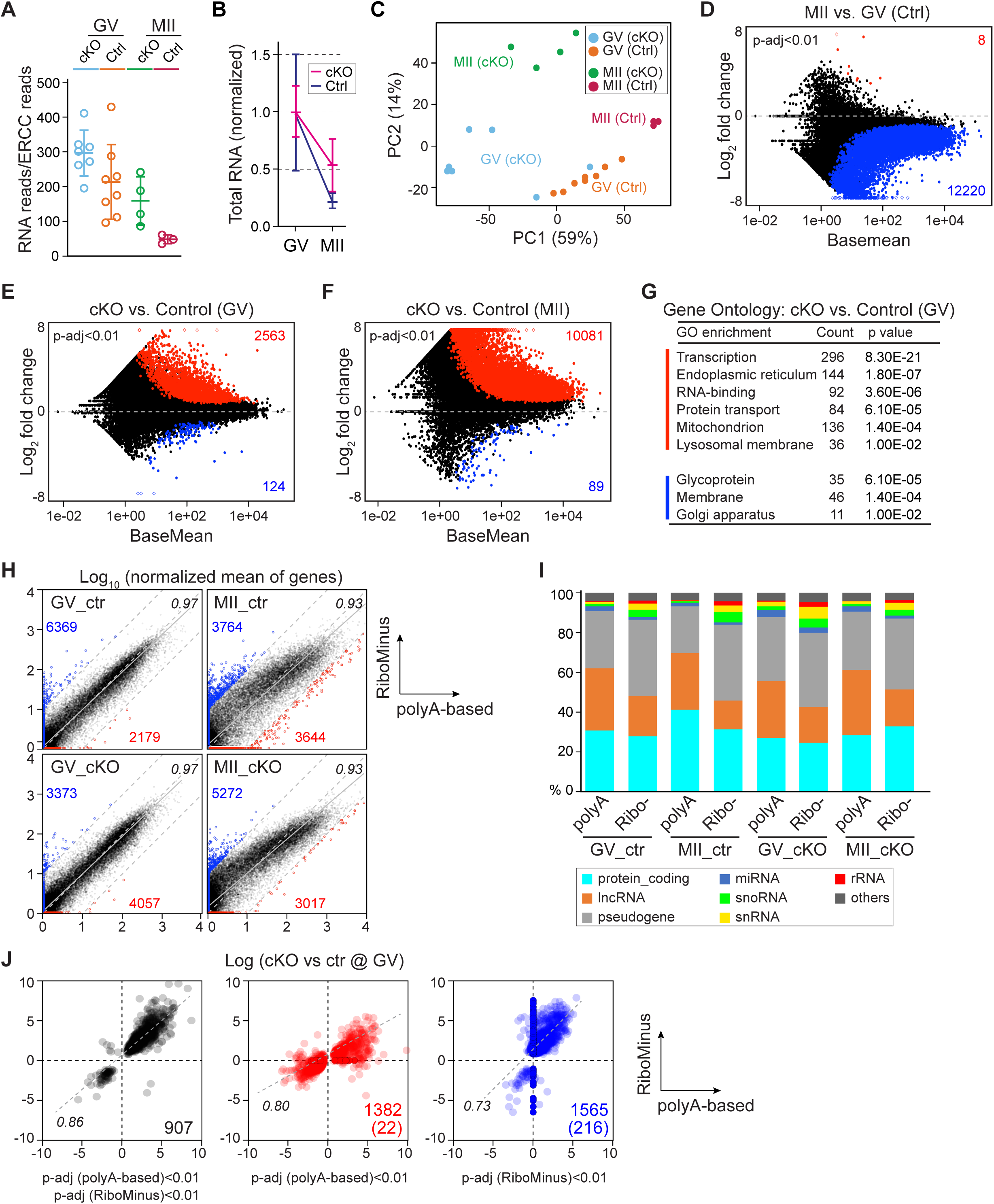
Non-poly(A) transcripts are also affected in *Exosc10*^*cKO*^ oocytes detected by RiboMinus RNA-seq. (**A**) Total RNA abundance indicated by RNA reads normalized to ERCC reads. (**B**) Further normalization of **A** by the mean value of the GV stage within each genotype. (**C**) Principal component analysis (PCA) of all libraries. Each dot represents one library, color coded by genotype and stage. (**D**) MA-plot of transcript changes from GV to MII in control oocytes. The increased and decreased transcripts are labeled by red and blue, respectively (both have *P*-adjust <0.01). (**E-F**) MA-plots of transcript changes in cKO vs. control oocytes at GV and MII stages. The increased and decreased transcripts are labeled by red and blue, respectively (*P*-adjust <0.01). (**G**) Gene ontology of **E**. Red: terms enriched by increased abundant transcripts; blue: terms enriched by decreased transcripts. (**H**) Plots of transcript abundance from poly(A) and RiboMinus sequencing. Normalized mean of genes: mean of genes counts from all ERCC-normalized libraries in each condition. In each plot, the x-axis is the log_10_ count value from poly(A) sequencing results and the y-axis is that from RiboMinus sequencing. Red dots/number, the outliers in poly(A) sequencing; blue dots/number, the outliers in RiboMinus sequencing. Three gray dashed lines: y=x, y=x+1 and y=x-1, which define the outliers in each sequencing result. Gray solid line: fitted line of all the dots. Black number: the correlation coefficient of each regression. (**I**) Gene type analysis in each condition. (**J**) Plots comparing the differentially accumulated transcripts from poly(A) and RiboMinus sequencing results. Black dots, transcripts with *P*-adjust <0.01 in both sequencing results; red dots, *P*-adjust <0.01 in poly(A)-based sequencing result; blue, *P*-adjust <0.01 in RiboMinus sequencing result. Numbers indicate transcripts shared by the two methods in each plot; number in parenthesis indicate transcripts detected only in one sequencing result.

In comparing the two RNA-seq approaches, we detected transcripts from 20,000 and 25,992 genes, with the poly(A)-based and RiboMinus methods, respectively. When aligning the normalized transcript counts, most detected transcripts exhibited a high correlation between the two methods in comparative conditions (0.97 for control and cKO group at the GV stage; 0.93 for control and cKO group at MII stage) (Figure 4H). However, we also identified poly(A)- preferred and RiboMinus-preferred transcripts which were either specifically detected by one method or having a deviation of more than 10-fold between the two methods (Figure 4H; Table S3). The poly(A) method covered more lncRNAs with potentially long poly(A) tails, and the RiboMinus method covered more snoRNA, snRNA and histone coding mRNAs that have short or no poly(A) tails, though the two methods covered similar levels of most protein coding transcripts, miRNA and pseudogene transcripts (Figure 4I). We also compared the differentially expressed transcript obtained from the two methods and found 907 transcripts significantly changed in both methods (correlation coefficient of 0.86) (Figure 4J). Additionally, 1,382 and 1,565 transcripts that were significantly changed in only one method also exhibited high correlation though not considered significantly changed from the other method (Figure 4J). From these observations, we conclude that both poly(A)-based and RiboMinus methods provide a comparable resolution in documenting changes in the transcriptome.

Additionally, though the cKO oocytes show normal percentage of read coverage in coding, UTR, intronic and intergenic regions, we analyzed the differentially expressed intronic and intergenic transcripts (Figure S5B) based on the RiboMinus RNA-seq data. We observed different abundance of intronic transcripts in cKO oocytes, which partly overlaps with coding transcripts (Figure S5C). However, the correlation between intronic and coding change was very low (0.50) (Figure S5D). Most of the intronic peaks were separate and of less intensity compared to the coding region peaks (Figure S5E), suggesting the intronic signal did not come from unsuccessful splicing, but are more likely to be unknown enhancer or noncoding transcripts. Similarly, separate peaks were also found in the intergenic transcripts (Figure S5F).

### cKO oocytes have disrupted endomembrane system

Nuclear envelope breakdown involves multiple components of the endomembrane system (36-38). Consistent with these earlier observations, the GO analysis of the cKO RNA-seq highlighted intracellular vesicle trafficking. Therefore, we examined a range of cytoplasmic vesicles including endosomes, ER, lysosomes and Golgi. RAB5-labeled early endosomes displayed abnormal aggregation and increased intensity in cKO oocytes (Figure 5A-B). In contrast, RAB7-labeled late endosomes were reduced (Figure 5C-D) suggesting insufficient endosome maturation (39). Live imaging of oocytes treated by ER-Tracker had a reduced signal, whereas the Lyso-Tracker treated oocytes had increased signal (Figure 5E).

**Figure 5.**
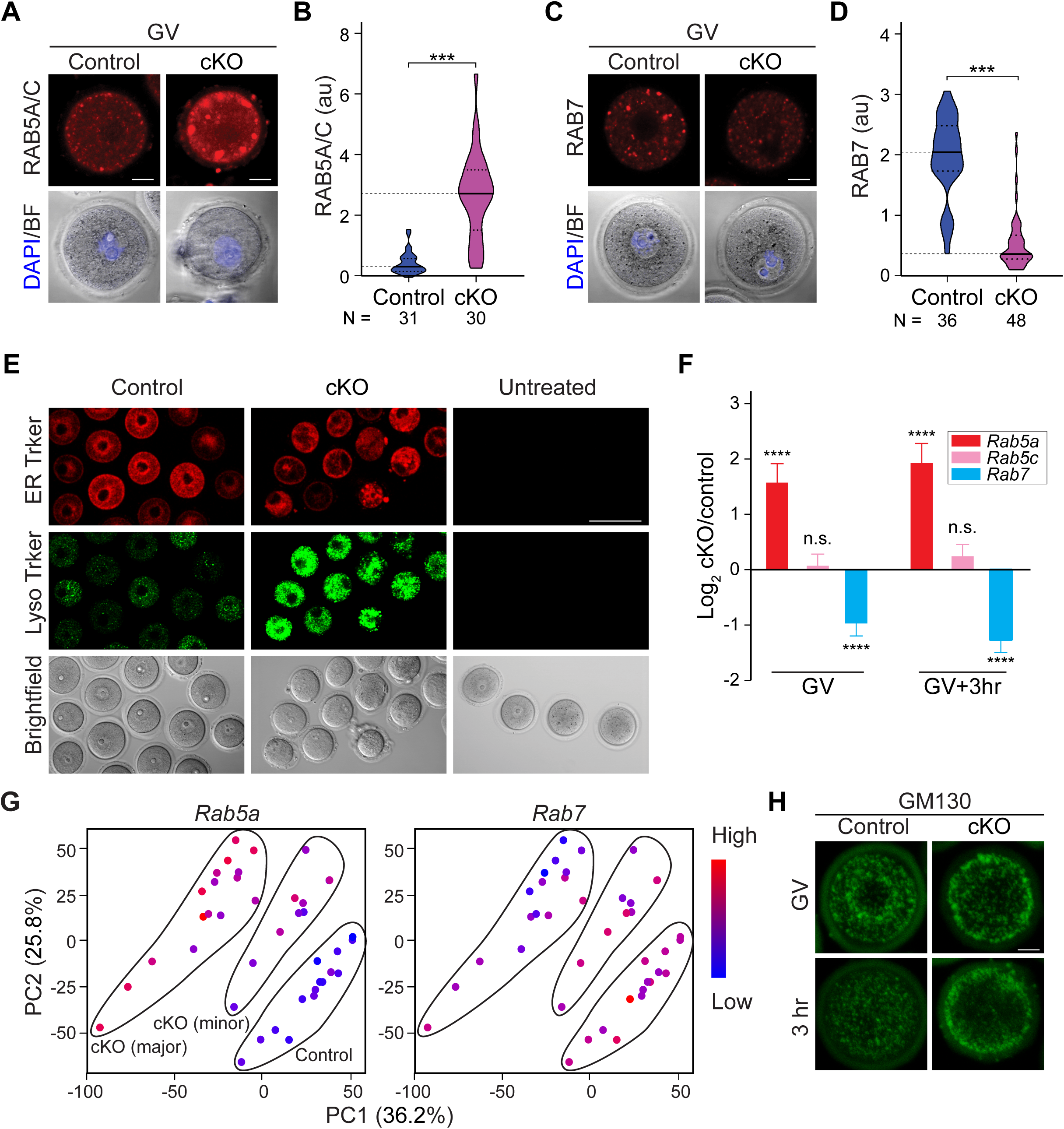
*Exosc10*^*cKO*^ oocytes have disrupted endomembrane system. (**A**) Immunostaining of oocyte RAB5A/C (early endosome) at the GV stage. (**B**) Quantification of RAB5A/C vesicle fluorescence intensity in **A**. (**C**) Same as **A**, but for RAB7 (late endosome). (**D**) Quantification of RAB7 vesicle intensities in **C**. For the violin plots in **B, D**, the inside horizontal lines inside represent the median and the dash lines represent the quartiles. **** *P* <0.0001, two-tailed Student’s t-test. Number of oocytes is indicated below each group. (**E**) Live imaging of oocytes derived from control and cKO incubated with ER Tracker and LysoTracker. The untreated group is in the right column. (**F**) A bar graph showing mean and standard error of log_2_ fold change of *Rab5A, Rab5C* and *Rab7* transcripts from single oocyte RNA-seq. **** *P* <0.0001, n.s. no significance, which are the *P*-adjust values by DESeq2 analysis. (**G**) PCA plots of *Rab5* (left) and *Rab7* (right) that is color-coded by the expression level of each transcript in individual oocytes. The control, cKO (major) and cKO (minor) groups are labeled in the *Rab5* plot. (**H**) Immunostaining of GM130 for Golgi apparatus at GV and 3 hr stages. Scale bars: 20 μm in **A, C, H**; 1 μm in **E**. Arbitrary fluorescence units (au) in **B, D**.

In the RNA-seq analyses, the abundance of *Rab5a* transcripts was significantly increased in cKO oocytes whereas *Rab7* transcripts were decreased (Figure 5F). The increased *Rab5a* and decreased *Rab7* transcript abundance in cKO (minor) and cKO (major) was confirmed by their expression levels on the PCA plot (Figure 5G). We tried to phenocopy the cKO oocytes by over-expressing *Rab5a-mVenus/Rab5c-mVenus* cRNA in GV oocytes. However, there was no obvious delay or arrest compared with the *mVenus* over-expression controls (Figure S6A-B). Nor did we observe defects in endosome maturation (Figure S6C-D), indicating that the endosome failure is likely to involve other impaired vesicles. We also detected the Golgi apparatus that normally exhibits a strong perinuclear signal and becomes accentuated after GVBD. However, the cKO oocytes showed a much stronger pericytoplasmic enrichment of Golgi and a reduced signal in the perinuclear region which remained unchanged after 3 hr culture (Figure 5H). In sum, formation of the endomembrane system was disrupted after EXOSC10 depletion.

### Inhibitory phosphorylated CDK1 persists in cKO oocytes to block lamina disassembly

The driving force of nuclear envelope breakdown is phosphorylation of nuclear lamina protein and nuclear pore components by active mitotic/meiotic kinases (40,41). To obtain better insight into GVBD, we divided oocytes into three groups based on lamin B integrity: intact, GVBD (early) and GVBD (late). The intact phase had an even and continuous distribution of lamin B surrounding the nucleus and was widely present among oocytes during 3 hr *ex vivo* culture. The GVBD (early) group emerged ∼1 hr, during which lamin B was still continuous (XY optical sections) but exhibiting much smaller enclosed areas and an unevenly shrunken pattern. Finally, after the nuclear envelope was breached by the spindle, lamin B became discontinuous and quickly disappeared, which defined the GVBD (late) phase. Lamin A/C staining precisely recapitulated that of lamin B (Figure S7A-D).

Given the well-studied control of CDK1 phosphorylation in meiosis, we hypothesized that the inhibitory phosphorylated CDK1 favors GV arrest while its elimination eventually results in GVBD. To demonstrate this, we explored the change of inhibitory phosphorylated CDK1 (p^T14/Y15^CDK1) during wildtype GVBD. In oocytes at the intact phase, the p^T14/Y15^CDK1 formed nuclear puncta. When entering the GVBD (early) phase, the puncta pattern underwent a constant decrease and eventually disappeared during the GVBD (late) phase (Figure 6A-B). We then hypothesized that the p^T14/Y15^CDK1 remained constant in cKO oocytes to block GVBD. At the GV stage, p^T14/Y15^CDK1 in cKO oocytes was similar to the control group. At GV3h when most control oocytes finished GVBD and exhibited a low level of p^T14/Y15^CDK1, the cKO oocytes retained p^T14/Y15^CDK1 puncta of high intensity (Figure 6C-D). Thus, failure in transiting CDK1 phosphorylation from inhibitory to active status results in the GV arrest in cKO oocytes.

**Figure 6.**
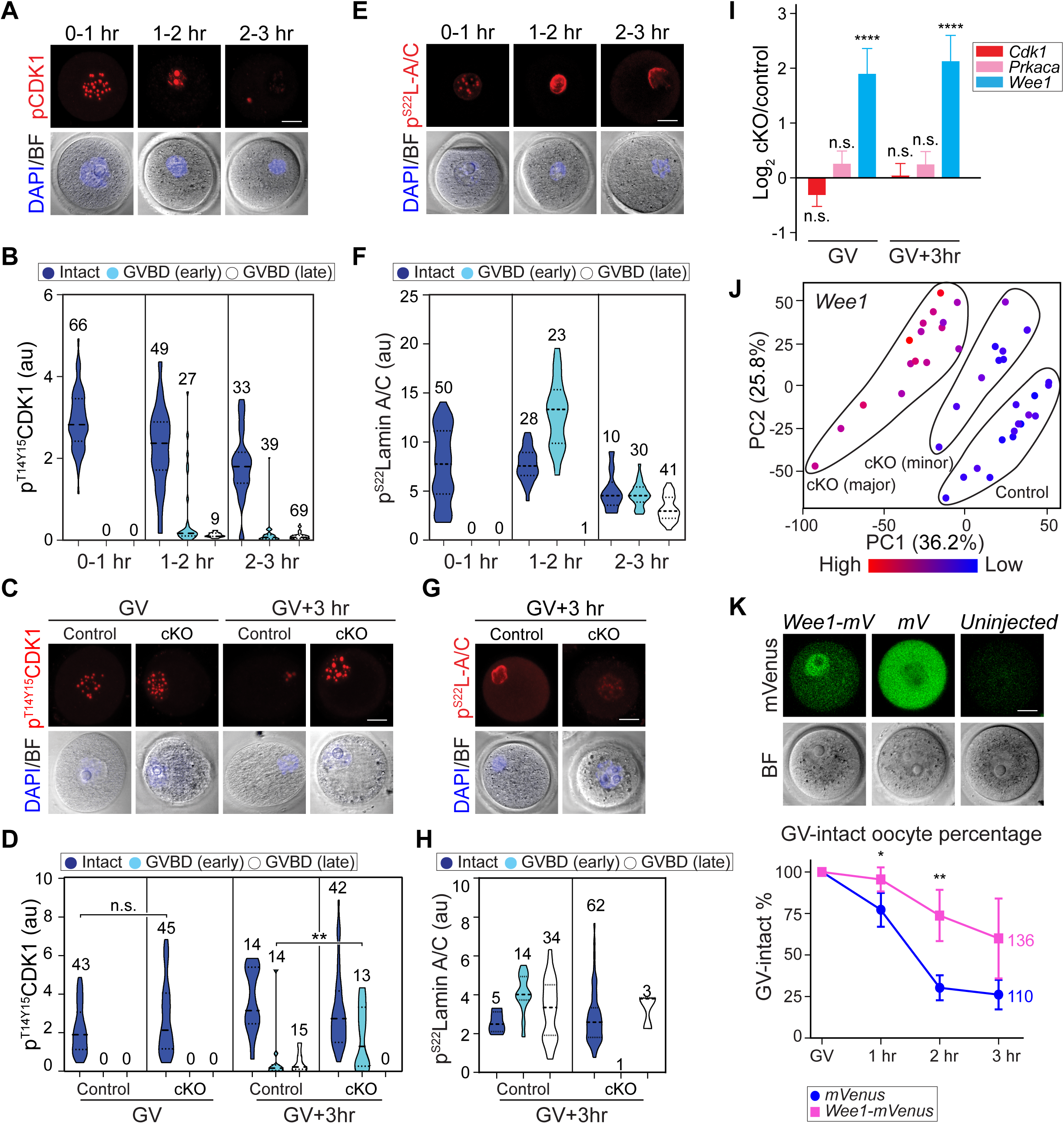
*Exosc10*^*cKO*^ oocytes with inhibitory CDK1 lack lamina phosphorylation and GVBD. **A**) Confocal fluorescence images of inhibitory phosphorylated CDK1 (p^T14/Y15^CDK1), bright-field and DAPI in wildtype oocytes at different time points during *ex vivo* culture and GVBD. Oocytes were collected every 30 min and combined into three stages: 0-1 hr includes 0 and 0.5 hr stages; 1-2 hr includes 1 and 1.5 hr stages; 2-3 hr includes 2, 2.5 and 3 hr stages. (**B**) Quantification of p^T14/Y15^CDK1 in **A**. Number of oocytes are indicated above each group. At each time points, oocytes are clustered as Intact, GVBD-early or GVBD-late based on lamin B integrity. (**C**) Representative images of p^T14/Y15^CDK1 in oocytes at GV and GV3h stages derived from control and cKO mice. (**D**) Quantification of p^T14/Y15^CDK1 fluorescence intensities in **C**. The numbers of oocytes from at least three experiments are indicated above each group. (**E**) Same as **A**, but for p^S22^lamin A/C (p^S22^L-A/C). (**F**) Same as **B**, but for p^S22^lamin A/C. (**G**) Same as **C**, but for p^S22^lamin A/C only at GV3h. (**H**) Same as **D**, but for p^S22^lamin A/C only at GV3h. (**I**) Bar graph of mean and standard error of log_2_ fold change of *Cdk1, Prkaca, Wee1* transcripts from the single oocyte RNA-seq. **** *P* <0.0001, n.s. no significance, which are the *P*-adjust values by DESeq2 analysis. (**J**) PCA plot of *Wee1* that is color-coded by the expression level in each oocyte. The control, cKO (major) and cKO (minor) groups are labeled in the plot. (**K**) Fluorescence of *Wee1-mVenus* and *mVenus* over-expression in GV-intact oocytes. The quantification of GV-intact oocyte percentage in 3 hr *ex vivo* culture is at the bottom. Each dot shows mean and standard deviation. ** *P* <0.01, * *P* <0.05 by two-tailed Student’s t-test. The horizontal lines in **B, D, F, H** represent the median and quartiles. Scale bars: 20 μm in **A, C, E, G, K**.

Next, we analyzed phosphorylation of lamin A/C (p^S22^lamin A/C) which is the direct substrate of active phosphorylated CDK1 (41). After 1 hr *ex vivo* culture, p^S22^lamin A/C displayed a nuclear puncta pattern in oocytes. Upon entering the GVBD (early) phase, p^S22^lamin A/C increased its abundance with concomitant loss of punctate loci and co-localized with lamin A/C. After progressing to the GVBD (late) phase, lamin A/C and p^S22^lamin A/C co-localized on the dissolving nuclear envelope albeit with decreased abundance (Figure 6E-F). These observations suggest that the activation of p^T14/Y15^CDK1 increases p^S22^lamin A/C, which results in nuclear envelope disassembly. In cKO oocytes, p^S22^lamin A/C failed to change from puncta to perinuclear localization at GV3h and failed to accumulate, both of which appear necessary for entering the GVBD (late) phase (Figure 6G-H).

The CDK1 phosphorylation could be regulated by upstream kinase and phosphorylase, including WEE1/2 and CDC25 (42,43). Our RNA-seq data documented that *Wee1* abundance was increased significantly in cKO oocytes (log_2_FC 1.92 and 2.16 at GV and GV3h, respectively, *P*-adjust <0.01), potentially contributing to the higher p^T14/Y15^CDK1 level (Figure 6I), while *Wee2* and *Cdc25* transcripts remained unchanged. The increased abundance of *Wee1* was confirmed by visualizing its expression level on the PCA plot (Figure 6J). To test whether the increase of *Wee1* may contribute to GVBD failure, we over-expressed *Wee1-mVenus* cRNA in wildtype GV oocytes. Nuclear expression of WEE1-mVenus was significantly increased in GV-intact oocytes within 3 hr *ex vivo* culture (Figure 6K) suggesting a delay and block of GVBD by WEE1 up-regulation. We also examined cAMP signaling which functions upstream of CDK1 phosphorylation to maintain the oocyte in a GV-intact state. However, cAMP levels and its known activators did not increase in cKO oocytes. These observations are consistent with signaling pathways downstream of cAMP facilitating persistent inhibitory CDK1 phosphorylation, resulting in lamin A/C phosphorylation and a block to GVBD (Figure S7E-F).

### EXOSC10 depletion results in pre-rRNA processing defects

rRNA undergoes dynamic processing and transport in oocytes (15). EXOSC10 is known to mediate 3’ rRNA processing (44). By sequencing rRNA at the GV stage and examining different genomic components of the rRNA (45), we observed an increased level in 5’ ETS and ITS1 regions, while unchanged levels of mature rRNA (including 18S, 5.8S and 28S), ITS2 and 3’ ETS regions (Figure 7A). When we looked further into the 3’ processing of the 18S, 28S and 5.8S rRNA, we found an increased ITS1 5’ end, which suggests defective 3’ end processing of pre-18S rRNA. However, the 5.8S and 28S 3’ ends don’t show obvious processing defects (Figure 7B). Given the known function of snoRNA in processing rRNA (44,46), we also examined snoRNA from our RiboMinus RNA-seq. 227 snoRNAs were detected and 10 exhibit significantly increased abundance, including 6 members of Snord family, namely *Snord80, Snord4a, Snord83b, Snord13*, *Gm24489* and *Snord14c*.

**Figure 7.**
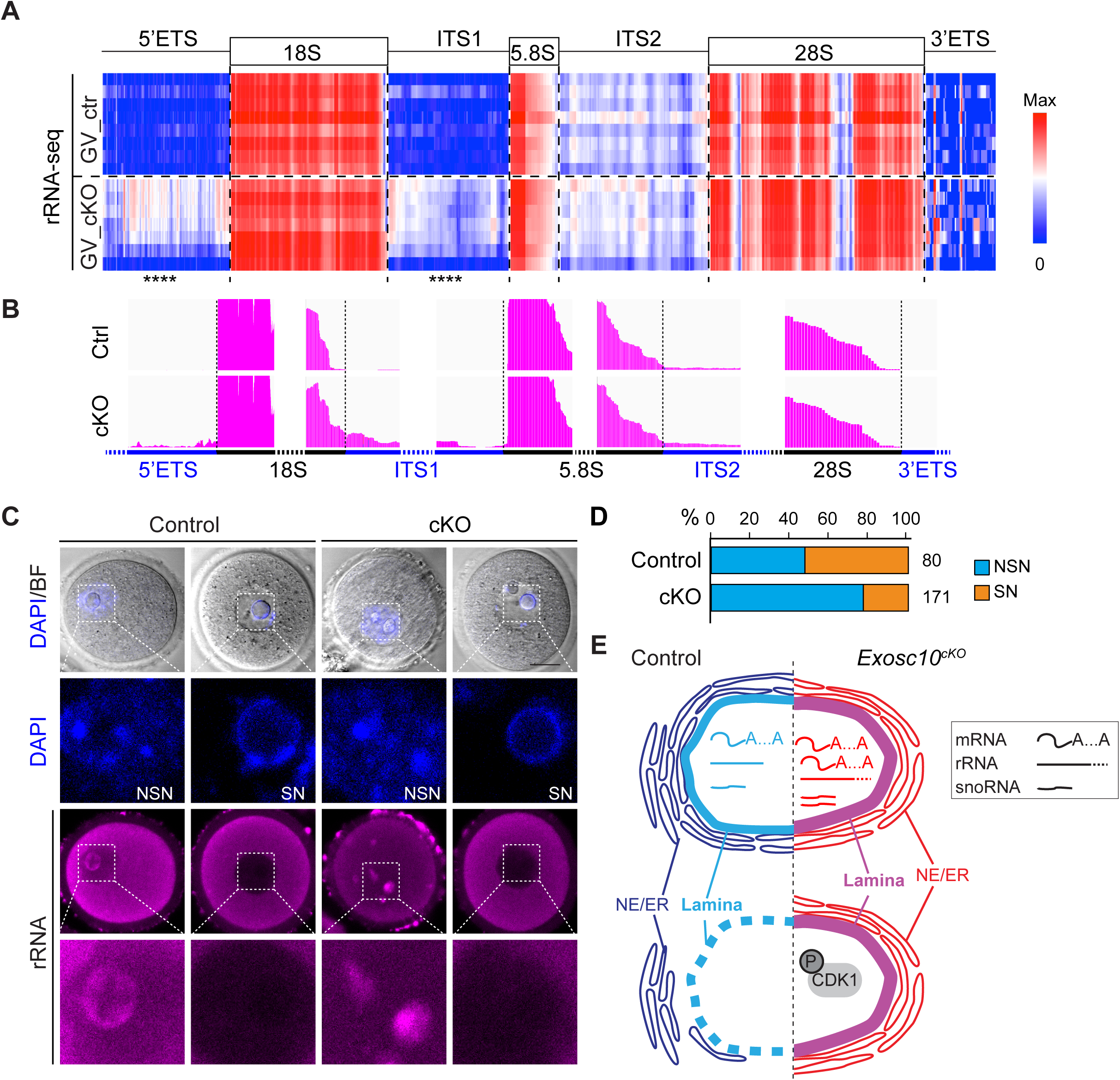
*Exosc10*^*cKO*^ oocytes have defective rRNA processing. (**A**) Heatmap of rRNA coverage obtained from GV stage rRNA sequencing mapped to the rRNA genomic region. **** *P*-adjust <0.0001 by DESeq2 differential analysis. (**B**) Integrated Genomics Viewer (IGV) graphs visualizing the boundary regions of rRNA. (**C**) rRNA immunostaining of GV oocytes. NSN, non-surrounded nucleolus; SN, surrounded nucleolus. Scale bar: 20 μm. (**D**) Bar graph showing percentage of NSN and SN oocytes of each genotype at the GV stage. (**E**) Working model of the characterized defects in *Exosc10*^*cKO*^ oocytes in which the absence of the RNase disrupts protein-coding genes level, rRNA processing and some snoRNA expression at the GV stage which prevents dephosphorylation of CDK1 and endomembrane trafficking. In the absence of this active cell cycle component, degradation of nuclear lamin B is delayed which preserves the nuclear envelope and defers GVBD causing decreased female fecundity.

We went on to localize rRNA by immunostaining with an anti-rRNA antibody. Interestingly, there was a high correlation between NSN (non-surrounded nucleolus) oocyte and the nuclear rRNA, and between SN (surrounded nucleolus) oocyte and the absent nuclear rRNA in both control and cKO groups. Moreover, the nucleolar rRNA pattern exists as puncta in the cKO oocytes rather than the peri-nucleolar pattern in the control oocytes (Figure 7C). Consistent with the heterogeneity of the oocytes and reduced GV progression, we observed relatively more NSN oocytes in the cKO group compared to the control (Figure 7D). Consequently, the overall translation activity of the oocyte decreased at the GV stage in cKO oocytes and became more severe at GV3h stage (Figure S7G). Thus, we conclude that rRNA processing is defective after EXOSC10 depletion.

When examining oocyte morphology, we noticed a modest reduction of cKO oocyte size (Figure S1I). To rule out the major growth defects, we compared our sequencing data to the published transcriptome of growing oocytes (GO) and fully-grown oocytes (FGO) (12). When analyzing the differentially expressed genes in cKO oocytes, we found most GO-specific transcripts remain unchanged (Figure S7H). Similarly, a very small percentage of developmentally differential genes from GO to FGO exhibit significant change in the cKO oocytes (76/826 GO-FGO increased and 30/590 GV-FGO decreased transcripts) (Figure S7I). The abundance of most transcripts was uniformly expressed without stage specificity. Thus, the subtle size decrease and the lack of GO-specific change confirmed that much of the growth process remains normal until the growth-to-maturation transition.

## DISCUSSION

Transcription and RNA degradation maintain a balance in sculpting the cellular transcriptome. During mouse oocyte maturation, the lack of transcription results in a more critical role of RNA degradation to shape the maternal transcriptome in preparation of zygotic control of the early embryo. Perturbation of RNA degradation in this process can result in disrupted meiosis or failed pre-implantation development. By evaluating the phenotype and RNA profile of conditional *Exosc10* knockout oocytes, we document important roles for EXOSC10 in forming a transcriptome necessary for endomembrane trafficking, meiotic cell cycle progression and processing rRNA (Figure 7E). The failure of EXOSC10 to sculpt the transcriptome of maturing oocytes causes a pronounced decline in female fertility.

To evaluate changes in the transcriptome of *Exosc10* knockout oocytes, we employed single oocyte RNA-seq using both poly(A)-dependent and poly(A)-independent methods. Different classes of RNAs, including protein-coding, lncRNA and snoRNA were dysregulated in the absence of EXOSC10. We compared the two methods to provide a window into how dynamic polyadenylation affects poly(A) based RNA-seq results. Basically, the transcript counts generated from the two methods exhibit similar resolution at the GV-intact stage. At the MII stage, most transcripts remain correlated, but there was a smaller coefficient and the poly(A) method was further enriched for dormant RNAs deadenylated during oocyte growth and highly polyadenylated after GVBD, including *Mos* and *Plat*. The poly(A)-bias is less significant in cKO MII eggs presumably due to observed defects in oocyte meiotic maturation (Figure 4H). Other than protein coding mRNA, the cKO oocytes also exhibit dysregulation of intronic and intergenic transcripts. Separate genome localization and discordant change from nearby protein coding genes suggest they could be potential, yet unknown, non-coding transcripts or truncated intermediates. Their increased abundance may result from either insufficient degradation by EXOSC10, or accumulation of abnormally processed transcripts. Exploring the function of these non-coding transcripts in oocyte maturation also may add insight to quality control of female germ cells.

Our study provides insight into the physiology of the oocyte growth-to-maturation transition. First, we document that EXOSC10 shapes the transcriptome encoding endomembrane components (endosomes, Golgi, ER, lysosomes, autophagosomes) which are known to actively participate in oocyte maturation (47-50). In control oocytes, we identify the initial reframing of the endomembrane system during GV to GV3h, including down-regulation of *Sec61g* and *Sec61b* which participate the NE-ER exchange (17). In EXOSC10 deficient oocytes, several Rab family member transcripts were more abundant and encode proteins known for intracellular membrane trafficking: *Rab5* for endosome maturation; *Rab34* for phagosome fusion with lysosome; as well as *Rab37* and *Atg16l2* for phagosome formation (51-53). We also observed similarities between the cKO Golgi and the diffuse Golgi pattern defined by an siRNA screen (54). By comparing the siRNA with our RNA-seq data, we identified several transcripts with decreased abundance in cKO oocytes (*Aurkb, Csnk2b, Nrbp, Camk1*) suggesting that abnormal Golgi in cKO oocytes could result from decreases in critical Golgi regulators. Future studies will be needed to address the role of different vesicles in ensuring oocyte quality.

Second, we show that EXOSC10 coordinates the transcriptome of CDK1/lamin pathway necessary for GVBD. It has been reported that active CDK1 phosphorylation is repressed by WEE2 and not WEE1 (43,55,56), due to the much higher expression level of WEE2. In our study, *Wee2* transcripts remain unchanged in the *Exosc10* cKO oocytes whereas *Wee1* becomes ∼4-fold more abundant. Overexpression of *Wee1* cRNA reduces the GVBD, which suggests that WEE1 could also repress CDK1 activation for resumption of meiosis I in the absence of timely degradation. On the other hand, the abundance of *Cdc25a/b/c* transcripts, which are known CDK1 activation phosphatases, remain largely unchanged (Table S1; Table S2). We also profiled the transcriptome at the MII stage which contains oocytes escaping from GV arrest and observe only partial or no defects during maturation. The differential abundance of transcripts could reflect another layer of EXOSC10 function following GVBD. However, due to possible secondary effects occurring throughout maturation (e.g., endomembrane and translation defects, inactive dormant RNAs, etc.), we did not pursue these investigations further.

Third, we demonstrate that EXOSC10 directly processes rRNA in oocytes. Through single oocyte rRNA sequencing, we document the association between rRNA processing and nuclear export that is reflected in NSN-to-SN progression. After EXOSC10 depletion, the failure of 5’ETS degradation and insufficient 18S rRNA 3’ trimming is associated with nuclear accumulation of rRNA, an increased percentage of NSN oocytes and eventually GV arrest. Moreover, rRNA defects contribute to the dysregulated transcriptome by disrupting protein translation which substantially impairs oocyte maturation. In addition to rRNA, the adenylation of some mRNAs may also be regulated by EXOSC10 (28). However, we did not observe any overall change in the poly(A) signal by FISH and the close correlation of differentially expressed genes using poly(A)-dependent and -independent methods supports minimal changes in poly-adenylation. Thus, we conclude that failure of rRNA processing and mRNA degradation play dominant roles in disrupting the transcriptome of cKO oocytes.

Lastly, our study provides insight into oocyte heterogeneity. Mutation of genes involved in oocyte RNA metabolism can result in infertility or subfertility despite similar decreases in the abundance of cognate transcripts. For example, infertility occurs when knocking-in an inactive allele of *Ago2* (*Ago2ADH*) (57), knocking out *Btg4* (4) and conditionally deleting *Dicer* (58), *Zfp36l2* (14), or *Tut4/7* (59). On the other hand, genetic ablation of an RNA binding complex member *Cnot6l* (60) and of *Exosc10* leads to subfertility and a variated oocyte phenotype. Potential explanations include different genes have different functions and substrates (*e*.*g*. siRNA, translation, uridylation) and affect fertility or viability by different mechanisms both direct and indirect. Secondly, there could be redundancies to compensate for the targeted loss. For EXOSC10, the redundant gene could be another RNA exosome-associated RNases, such as DIS3 (44). Thirdly, the heterogeneity could come from stochastic variations in oocyte development. For example, chromosome condensation may happen at various time in a non-specific order, resulting in different nuclear configurations and residual transcription activity. Fourthly, conditional knockout mice may vary slightly in temporal and spatial expression of *Zp3-cre* causing variated defects.

In conclusion, our data document transcriptome remodeling in the transition from oocyte growth to maturation. Combining phenotypic characterization and sequence analysis, we conclude that the EXOSC10-depleted GV oocytes have less potential to organize germinal vesicle breakdown because of defects in endomembrane trafficking, CDK1 phosphorylation activation and rRNA maturation which collectively disrupts resumption of meiosis I. These results are consistent with a growing body of experimental data that RNA degradation governs the later stages of oogenesis while placing the oocyte’s gain of developmental potential into a more refined time window.

## DATA AVAILABILITY

The sequencing data reported in this study has been deposited in the Gene Expression Omnibus website with accession code GSE141190.

## SUPPLEMENTARY DATA

Supplementary Data are available at NAR Online.

## ACKNOWLEDGEMENTS

We appreciate the critical reading of the manuscript by Shaohe Wang, Ph.D. and Yangu Zhao, Ph.D. as well as useful discussions with all the members of the Jurrien Dean lab. We appreciate the bioinformatic training and discussion from Cameron Palmer, Ph.D. D.W. and J.D conceived the project. D.W. performed the experiments and bioinformatic analysis. D.W. and J.D. wrote manuscript.

## REFERENCE

1. Jose-Miller, A.B., Boyden, J.W. and Frey, K.A. (2007) Infertility. Am. Fam. Physician, 75, 849–856.

2. Macklon, N.S., Geraedts, J.P. and Fauser, B.C. (2002) Conception to ongoing pregnancy: the ‘black box’ of early pregnancy loss. Hum. Reprod. Update, 8, 333–343.

3. Paynton, B.V., Rempel, R. and Bachvarova, R. (1988) Changes in state of adenylation and time course of degradation of maternal mRNAs during oocyte maturation and early embryonic development in the mouse. Dev. Biol., 129, 304–314.

4. Yu, C., Ji, S.Y., Sha, Q.Q., Dang, Y., Zhou, J.J., Zhang, Y.L., Liu, Y., Wang, Z.W., Hu, B., Sun, Q.Y. et al. (2016) BTG4 is a meiotic cell cycle-coupled maternal-zygotic-transition licensing factor in oocytes. Nat. Struct. Mol. Biol., 23, 387–394.

5. Ma, J., Fukuda, Y. and Schultz, R.M. (2015) Mobilization of dormant Cnot7 mRNA promotes deadenylation of maternal transcripts during mouse oocyte maturation. Biol. Reprod., 93, 48.

6. Svoboda, P., Stein, P., Hayashi, H. and Schultz, R.M. (2000) Selective reduction of dormant maternal mRNAs in mouse oocytes by RNA interference. Development, 127, 4147–4156.

7. Strickland, S., Huarte, J., Belin, D., Vassalli, A., Rickles, R.J. and Vassalli, J.D. (1988) Antisense RNA directed against the 3’ noncoding region prevents dormant mRNA activation in mouse oocytes. Science, 241, 680–684.

8. Metchat, A., Akerfelt, M., Bierkamp, C., Delsinne, V., Sistonen, L., Alexandre, H. and Christians, E.S. (2009) Mammalian heat shock factor 1 is essential for oocyte meiosis and directly regulates Hsp90alpha expression. J. Biol. Chem., 284, 9521–9528.

9. Burns, K.H., Viveiros, M.M., Ren, Y., Wang, P., DeMayo, F.J., Frail, D.E., Eppig, J.J. and Matzuk, M.M. (2003) Roles of NPM2 in chromatin and nucleolar organization in oocytes and embryos. Science, 300, 633–636.

10. Wasielak, M., Wiesak, T., Bogacka, I., Jalali, B.M. and Bogacki, M. (2016) Zygote arrest 1, nucleoplasmin 2, and developmentally associated protein 3 mRNA profiles throughout porcine embryo development in vitro. Theriogenology, 86, 2254–2262.

11. Schultz, R.M., Stein, P. and Svoboda, P. (2018) The oocyte-to-embryo transition in mouse: past, present, and future. Biol. Reprod., 99, 160–174.

12. Veselovska, L., Smallwood, S.A., Saadeh, H., Stewart, K.R., Krueger, F., Maupetit-Mehouas, S., Arnaud, P., Tomizawa, S., Andrews, S. and Kelsey, G. (2015) Deep sequencing and de novo assembly of the mouse oocyte transcriptome define the contribution of transcription to the DNA methylation landscape. Genome Biol., 16, 209.

13. Kageyama, S., Liu, H., Kaneko, N., Ooga, M., Nagata, M. and Aoki, F. (2007) Alterations in epigenetic modifications during oocyte growth in mice. Reproduction, 133, 85–94.

14. Dumdie, J.N., Cho, K., Ramaiah, M., Skarbrevik, D., Mora-Castilla, S., Stumpo, D.J., Lykke-Andersen, J., Laurent, L.C., Blackshear, P.J., Wilkinson, M.F. et al. (2018) Chromatin modification and global transcriptional silencing in the oocyte mediated by the mRNA decay activator ZFP36L2. Dev. Cell, 44, 392–402 e397.

15. Shishova, K.V., Lavrentyeva, E.A., Dobrucki, J.W. and Zatsepina, O.V. (2015) Nucleolus-like bodies of fully-grown mouse oocytes contain key nucleolar proteins but are impoverished for rRNA. Dev. Biol., 397, 267–281.

16. Kishimoto, T. (2018) MPF-based meiotic cell cycle control: Half a century of lessons from starfish oocytes. Proc. Jpn. Acad. Ser. B. Phys. Biol. Sci., 94, 180–203.

17. Anderson, D.J. and Hetzer, M.W. (2008) Reshaping of the endoplasmic reticulum limits the rate for nuclear envelope formation. J. Cell Biol., 182, 911–924.

18. Liu, J., Prunuske, A.J., Fager, A.M. and Ullman, K.S. (2003) The COPI complex functions in nuclear envelope breakdown and is recruited by the nucleoporin Nup153. Dev. Cell, 5, 487–498.

19. Hartley, J.L., Zachos, N.C., Dawood, B., Donowitz, M., Forman, J., Pollitt, R.J., Morgan, N.V., Tee, L., Gissen, P., Kahr, W.H.A. et al. (2010) Mutations in TTC37 cause Trichohepatoenteric Syndrome (Phenotypic Diarrhea of Infancy). Gastroenterology, 138, 2388–2398.

20. Rudnik-Schöneborn, S., Senderek, J., Jen, J.C., Houge, G., Seeman, P., Puchmajerová, A., Graul-Neumann, L., Seidel, U., Korinthenberg, R., Kirschner, J. et al. (2013) Pontocerebellar hypoplasia type 1: Clinical spectrum and relevance of EXOSC3 mutations. Neurology, 80, 438–446.

21. Weissbach, S., Langer, C., Puppe, B., Nedeva, T., Bach, E., Kull, M., Bargou, R., Einsele, H., Rosenwald, A., Knop, S. et al. (2015) The molecular spectrum and clinical impact of DIS3 mutations in multiple myeloma. Br. J. Haematol., 169, 57–70.

22. Carneiro, T., Carvalho, C., Braga, J., Rino, J., Milligan, L., Tollervey, D. and Carmo-Fonseca, M. (2007) Depletion of the yeast nuclear exosome subunit Rrp6 results in accumulation of polyadenylated RNAs in a discrete domain within the nucleolus. Mol. Cell Biol., 27, 4157–4165.

23. Rolfsmeier, M.L., Laughery, M.F. and Haseltine, C.A. (2011) Repair of DNA double-strand breaks induced by ionizing radiation damage correlates with upregulation of homologous recombination genes in Sulfolobus solfataricus. J. Mol. Biol., 414, 485–498.

24. Domingo-Prim, J., Endara-Coll, M., Bonath, F., Jimeno, S., Prados-Carvajal, R., Friedlander, M.R., Huertas, P. and Visa, N. (2019) EXOSC10 is required for RPA assembly and controlled DNA end resection at DNA double-strand breaks. Nat. Commun., 10, 2135.

25. van Dijk, E.L., Schilders, G. and Pruijn, G.J. (2007) Human cell growth requires a functional cytoplasmic exosome, which is involved in various mRNA decay pathways. Rna, 13, 1027–1035.

26. Knight, J.R.P., Bastide, A., Peretti, D., Roobol, A., Roobol, J., Mallucci, G.R., Smales, C.M. and Willis, A.E. (2016) Cooling-induced SUMOylation of EXOSC10 down-regulates ribosome biogenesis. Rna, 22, 623–635.

27. Pefanis, E., Wang, J., Rothschild, G., Lim, J., Kazadi, D., Sun, J., Federation, A., Chao, J., Elliott, O., Liu, Z.P. et al. (2015) RNA exosome-regulated long non-coding RNA transcription controls super-enhancer activity. Cell, 161, 774–789.

28. West, S., Gromak, N., Norbury, C.J. and Proudfoot, N.J. (2006) Adenylation and exosome-mediated degradation of cotranscriptionally cleaved pre-messenger RNA in human cells. Mol. Cell, 21, 437–443.

29. Jamin, S.P., Petit, F.G., Kervarrec, C., Smagulova, F., Illner, D., Scherthan, H. and Primig, M. (2017) EXOSC10/Rrp6 is post-translationally regulated in male germ cells and controls the onset of spermatogenesis. Sci. Rep., 7, 15065.

30. Park, S.J., Shirahige, K., Ohsugi, M. and Nakai, K. (2015) DBTMEE: a database of transcriptome in mouse early embryos. Nucleic Acids Res., 43, D771–776.

31. Lewandoski, M., Wassarman, K.M. and Martin, G.R. (1997) Zp3-cre, a transgenic mouse line for the activation or inactivation of loxP-flanked target genes specifically in the female germ line. Curr. Biol., 7, 148–151.

32. Macaulay, I.C., Teng, M.J., Haerty, W., Kumar, P., Ponting, C.P. and Voet, T. (2016) Separation and parallel sequencing of the genomes and transcriptomes of single cells using G&T-seq. Nat. Protoc., 11, 2081–2103.

33. Lemieux, C., Marguerat, S., Lafontaine, J., Barbezier, N., Bahler, J. and Bachand, F. (2011) A Pre-mRNA degradation pathway that selectively targets intron-containing genes requires the nuclear poly(A)-binding protein. Mol. Cell, 44, 108–119.

34. Tseng, C.K., Wang, H.F., Schroeder, M.R. and Baumann, P. (2018) The H/ACA complex disrupts triplex in hTR precursor to permit processing by RRP6 and PARN. Nat. Commun., 9, 5430.

35. Guttinger, S., Laurell, E. and Kutay, U. (2009) Orchestrating nuclear envelope disassembly and reassembly during mitosis. Nat. Rev. Mol. Cell Biol., 10, 178–191.

36. Audhya, A., Desai, A. and Oegema, K. (2007) A role for Rab5 in structuring the endoplasmic reticulum. J. Cell Biol., 178, 43–56.

37. Chung, G.H.C., Domart, M.C., Peddie, C., Mantell, J., McLaverty, K., Arabiotorre, A., Hodgson, L., Byrne, R.D., Verkade, P., Arkill, K. et al. (2018) Acute depletion of diacylglycerol from the cis-Golgi affects localized nuclear envelope morphology during mitosis. J. Lipid Res., 59, 1402–1413.

38. Ivanov, A., Pawlikowski, J., Manoharan, I., van Tuyn, J., Nelson, D.M., Rai, T.S., Shah, P.P., Hewitt, G., Korolchuk, V.I., Passos, J.F. et al. (2013) Lysosome-mediated processing of chromatin in senescence. J. Cell Biol., 202, 129–143.

39. Poteryaev, D., Datta, S., Ackema, K., Zerial, M. and Spang, A. (2010) Identification of the switch in early-to-late endosome transition. Cell, 141, 497–508.

40. Peter, M., Nakagawa, J., Doree, M., Labbe, J.C. and Nigg, E.A. (1990) In vitro disassembly of the nuclear lamina and M phase-specific phosphorylation of lamins by cdc2 kinase. Cell, 61, 591–602.

41. Heald, R. and McKeon, F. (1990) Mutations of phosphorylation sites in lamin A that prevent nuclear lamina disassembly in mitosis. Cell, 61, 579–589.

42. Han, S.J. and Conti, M. (2006) New pathways from PKA to the Cdc2/cyclin B complex in oocytes: Wee1B as a potential PKA substrate. Cell Cycle, 5, 227–231.

43. Oh, J.S., Han, S.J. and Conti, M. (2010) Wee1B, Myt1, and Cdc25 function in distinct compartments of the mouse oocyte to control meiotic resumption. J. Cell Biol., 188, 199–207.

44. Davidson, L., Francis, L., Cordiner, R.A., Eaton, J.D., Estell, C., Macias, S., Caceres, J.F. and West, S. (2019) Rapid depletion of DIS3, EXOSC10, or XRN2 reveals the immediate impact of exoribonucleolysis on nuclear RNA metabolism and transcriptional control. Cell Rep., 26, 2779–2791 e2775.

45. Grozdanov, P., Georgiev, O. and Karagyozov, L. (2003) Complete sequence of the 45-kb mouse ribosomal DNA repeat: analysis of the intergenic spacer. Genomics, 82, 637–643.

46. Kiss, T. (2002) Small nucleolar RNAs: an abundant group of noncoding RNAs with diverse cellular functions. Cell, 109, 145–148.

47. Lowther, K.M., Nikolaev, V.O. and Mehlmann, L.M. (2011) Endocytosis in the mouse oocyte and its contribution to cAMP signaling during meiotic arrest. Reproduction, 141, 737–747.

48. Moreno, R.D., Schatten, G. and Ramalho-Santos, J. (2002) Golgi apparatus dynamics during mouse oocyte in vitro maturation: effect of the membrane trafficking inhibitor brefeldin A. Biol. Reprod., 66, 1259–1266.

49. FitzHarris, G., Marangos, P. and Carroll, J. (2007) Changes in endoplasmic reticulum structure during mouse oocyte maturation are controlled by the cytoskeleton and cytoplasmic dynein. Dev. Biol., 305, 133–144.

50. Shin, S.W., Vogt, E.J., Jimenez-Movilla, M., Baibakov, B. and Dean, J. (2017) Cytoplasmic cleavage of DPPA3 is required for intracellular trafficking and cleavage-stage development in mice. Nat. Commun., 8, 1643.

51. Sheng, Y., Song, Y., Li, Z., Wang, Y., Lin, H., Cheng, H. and Zhou, R. (2018) RAB37 interacts directly with ATG5 and promotes autophagosome formation via regulating ATG5-12-16 complex assembly. Cell Death Differ., 25, 918–934.

52. Ishibashi, K., Fujita, N., Kanno, E., Omori, H., Yoshimori, T., Itoh, T. and Fukuda, M. (2011) Atg16L2, a novel isoform of mammalian Atg16L that is not essential for canonical autophagy despite forming an Atg12-5-16L2 complex. Autophagy, 7, 1500–1513.

53. Kasmapour, B., Gronow, A., Bleck, C.K., Hong, W. and Gutierrez, M.G. (2012) Size-dependent mechanism of cargo sorting during lysosome-phagosome fusion is controlled by Rab34. Proc. Natl. Acad. Sci. U.S.A., 109, 20485–20490.

54. Chia, J., Goh, G., Racine, V., Ng, S., Kumar, P. and Bard, F. (2012) RNAi screening reveals a large signaling network controlling the Golgi apparatus in human cells. Mol. Syst. Biol., 8, 629.

55. Han, S.J., Chen, R., Paronetto, M.P. and Conti, M. (2005) Wee1B is an oocyte-specific kinase involved in the control of meiotic arrest in the mouse. Curr. Biol., 15, 1670–1676.

56. Mitra, J. and Schultz, R.M. (1996) Regulation of the acquisition of meiotic competence in the mouse: changes in the subcellular localization of cdc2, cyclin B1, cdc25C and wee1, and in the concentration of these proteins and their transcripts. J. Cell Sci., 109 (Pt 9), 2407–2415.

57. Stein, P., Rozhkov, N.V., Li, F., Cardenas, F.L., Davydenko, O., Vandivier, L.E., Gregory, B.D., Hannon, G.J. and Schultz, R.M. (2015) Essential Role for endogenous siRNAs during meiosis in mouse oocytes. PLoS Genet., 11, e1005013.

58. Murchison, E.P., Stein, P., Xuan, Z., Pan, H., Zhang, M.Q., Schultz, R.M. and Hannon, G.J. (2007) Critical roles for Dicer in the female germline. Genes Dev., 21, 682–693.

59. Morgan, M., Much, C., DiGiacomo, M., Azzi, C., Ivanova, I., Vitsios, D.M., Pistolic, J., Collier, P., Moreira, P.N., Benes, V. et al. (2017) mRNA 3’ uridylation and poly(A) tail length sculpt the mammalian maternal transcriptome. Nature, 548, 347–351.

60. Sha, Q.Q., Yu, J.L., Guo, J.X., Dai, X.X., Jiang, J.C., Zhang, Y.L., Yu, C., Ji, S.Y., Jiang, Y., Zhang, S.Y. et al. (2018) CNOT6L couples the selective degradation of maternal transcripts to meiotic cell cycle progression in mouse oocyte. EMBO J., 37.

