## Supplementary Figures for "EXOSC10 sculpts the transcriptome during the growth-to-maturation transition in mouse oocytes"

#### SUPPLEMANTERY FIGURES AND TABLES

**Figure S1. Characterization of *Exosc10* oocyte-conditional knockout mice.** (**A**) Schematic of the strategy to generate *Exosc10* knockout alleles using CRISPR/Cas9 in which two gRNAs were used to delete exons 4-10. (**B**) Schematic of mRNAs generated from the floxed and cKO alleles. In the latter, exons 4-10 are absent, and an early termination occurs in exon 11. (**C**) Number of pups with indicated genotypes. No *Exosc10* homozygous null pups survived to term. (**D**) qRT-PCR of *Exosc10* in single oocytes obtained from controls and cKO females. Left panel: single oocytes from GV stage onward; right panel: growing oocytes isolated from 10-day females. Two primer pairs were used (see **B)**. Primer pair 1, shown here, detects the region deleted after loxP recombination; primer pair 2 detects a region shared by the mRNAs transcribed from the *Exosc10* floxed and cKO alleles. (**E**) Representative ovaries (12 weeks old) dissected from control and cKO mice. (**F**) Ovary/body weight ratio of control and cKO females (12 weeks old). Number of ovaries is indicated below each group. (**G**) Histology of periodic acid-Schiff stained ovaries. (**H**) Quantification of fully-grown oocytes in single (2 $\mu$m) sections from **G**. Number of ovaries is indicated below each group. (**I**) Diameter of control and cKO oocytes. The horizontal lines represent the median and quartiles. (**J**) *Ex vivo* maturation of control and cKO GV oocytes through meiosis (MI, MII). The number of oocytes is indicated below each group. (**K**) Number of ovulated eggs collected from control and cKO female oviducts after hormonal stimulation with eCG and hCG. The number of females is indicated below each group. (**L**) Immunostaining of microtubule-organizing center (MTOC) proteins pericentrin (PCNT) and $\gamma$-tubulin, indicated by arrowheads. (**M**) Confocal fluorescence and bright-field images of EXOSC10-mVenus localization during control oocyte maturation and pre-implantation development. (**N**) *Ex vivo* culture of pre-implantation development of the embryos derived from control and cKO female mice. Bright-field images were obtained at E1.5, E2.5 and E3.5. (**O**) Quantification of embryonic progression at each time point in **N**. In **F**, **H**, **K**, the horizontal lines represent the mean and standard deviation. Scale bars: 1 mm in **E**, **G**; 20 $\mu$m in **L**, **M**; 100 $\mu$m in **N**. **** *P* <0.0001, ** *P* <0.05, two-tailed Student's t-test.

**Figure S2. EXOSC10 degrades poly(A) RNA in oocytes.** (**A, C, E**) Confocal fluorescence images showing poly(A) RNA FISH, bright-field and DAPI of wildtype oocytes at GV, GV+3hr (GV3h) and MII stages in **A**, of oocytes over expressing *mVenus*, *Exosc10-mVenus or dExosc10-mVenus* cRNA at GV, GV3h stages in **C**, and of oocytes derived from controls and cKO females at GV, GV3h in **E**. The oocytes at GV3h stage were obtained by *ex vivo* culture of GV oocytes in maturation medium for 3 hr. Note that the cytoplasmic EXOSC10-mVenus is due to the protein expansion to the cytoplasm after GVBD. (**B, D, F**). Quantification of overall poly(A) RNA fluorescence of **A**, **C**, **E**, respectively. The horizontal lines represent the mean and standard deviation. In **B**, Nc1, negative control 1, without probe incubation; Nc2, negative control 2, treated by RNase A before probe incubation. Number of oocytes indicated below each group. **** *P* <0.0001, * *P* <0.05 in **D** by one-way ANOVA. Scale bars: 20 $\mu$m in **A**, **C**, **E**.

**Figure S3. Validation of linear amplification of poly(A) RNA-seq libraries by ERCC RNA spike-in mix.** Each scatter plot represents the linear regression of ERCC reads with the original concentration. The red number is the covariant of coefficient and the blue number is the ERCC components (92 in total) detected. Libraries are from GV_ctr, control GV oocytes; GV.3_ctr, control GV oocytes incubated for 3 hr; MII_ctr, control MII oocytes; GV_cKO, cKO GV oocytes; GV.3_cKO, cKO GV oocyte incubated for 3 hr; MII_cKO, cKO MII oocytes.

**Figure S4. Validation of linear amplification of RiboMinus RNA-seq libraries by ERCC RNA spike-in mix.** Each scatter plot represents the linear regression of ERCC reads with the original concentration. The red number is the covariant of coefficient and the blue number is the ERCC components (92 in total) detected.

**Figure S5. Intronic and intergenic RNAs change by EXOSC10 depletion.** (**A**) Example plots showing the read coverage of more abundant genes in cKO by RiboMinus RNA-seq. (**B**) Picard analysis of poly(A) RNA-seq and RiboMinus RNA-seq to map the reads on coding, UTR, intronic, intergenic regions. (**C**) Bar graph showing the differentially expressed intronic genes compared to the coding genes. There are some overlapping transcripts that change coordinately for coding and intronic sequences. (**D**) Correlation of intronic transcripts and coding regions of the same gene ID. (**E**) Examples of differentially expressed coding and intronic regions of the same genes. Red/blue arrows indicate more and less abundant intronic peaks, respectively. (**F**) Examples of differentially expressed intergenic regions and the surrounding coding genes. Red/blue arrows indicate more and less abundant intergenic peaks, respectively.

**Figure S6. Over-expression of *Rab5a* and *Rab5c* cRNA failed to phenocopy *Exosc10$^{cKO}$* oocytes.** (**A**) Brightfield images after *ex vivo* culture (0, 3 hr, 20 hr) of wildtype oocytes injected with cRNA of *Rab5a-mVenus*, *Rab5c-mVenus*, *Rab5a-mVenus/Rab5c-mVenus* and *mVenus*. (**B**) Bar graphs of oocyte maturation from GVBD (BD) to MII in **A** in two independent experiments. The total number of oocytes is indicated above each bar graph. (**C-D**) Fluorescent images of oocytes after co-injection of cRNA of *Rab5a-mVenus* and *Rab5c-mVenus* which failed to decrease RAB7. Scale bars: 100 μm in **A**, **C, D**.

**Figure S7. Phenotype analysis of *Exosc10$^{cKO}$* oocytes.** (**A, C**) Representative figures of immunostaining of lamin B and lamin A/C at 0-1 hr, 1-2 hr and 2-3 hr of wildtype GV oocytes during *ex vivo* culture, which defines Intact, GVBD (early) and GVBD (late) oocytes in this study. (**B, D**) Quantification of lamin B and lamin A/C. The horizontal lines represent the median and quartiles. (**E-F**) cAMP level does not increase in *Exosc10$^{cKO}$* oocytes. (**E**) Confocal fluorescence images of cAMP in control and Exosc10 cKO GV oocytes. Scale bars: 20 μm. (**F**) A bar graph of the mean and standard error of $\log_2$ fold change of *Gpr12* (G protein-coupled receptor 12), *Gpr3* (G protein-coupled receptor 3), *Pde3a* (phosphodiesterase 3A), *Adcy3* (adenylate cyclase 3), *Adcy9* (adenylate cyclase 9) from single oocyte RNA-seq data sets. **** $P$ <0.0001, *** $P$ <0.001, ** $P$ <0.01, * $P$ <0.05, n.s. no significance, which are the *P*-adjust values in the DESeq2 analysis. (**G**) Translation activity detection by HPG labeling at GV and GV3h stages. (**H-I**) Transcriptome comparison between differentially expressed genes in *Exosc10$^{cKO}$* oocytes and the growth stage datasets. (**H**). A bar graph showing differentially expressed overlapping transcripts in GO (growing oocytes) and FGO (fully grown oocytes) from GSE70116. (**I**) MA plot mapping accumulated and degraded transcripts in GO-FGO *Exosc10$^{cKO}$* oocytes.

**Table S1.** Differential expression analysis of the single oocyte poly(A)-based RNA-seq by DESeq2, including GV_cKO vs GV_ctr, GV3h_cKO vs GV3h_ctr, MII_cKO vs MII_ctr, GV3h_ctr vs GV_ctr, MII_ctr vs GV3h_ctr, and cKO_major vs cKO_minor. Each differential analysis result contains the $\log_2$ fold change, the standard error, the *P*-adjust values of the significantly changed transcripts (*P*-adjust < 0.01) and gene symbols.

**Table S2.** Differential expression analysis of the single oocyte RiboMinus RNA-seq by DESeq2, including GV_cKO vs GV_ctr, MII_cKO vs MII_ctr, and MII_ctr vs GV_ctr. Each differential analysis result contains the $\log_2$ fold change, the standard error and the *P*-adjust values of the significantly changed transcripts (*P*-adjust < 0.01) and gene symbols.

**Table S3.** Significantly deviated transcripts sequenced from poly(A) RNA-seq and RiboMinus RNA-seq at each stage/genotype. Each sheet contains gene ID, normalized express level of all the sequenced oocytes within the same treatment, and gene symbols. The genes are either detected only by one method or exhibit a difference of more than 10-fold (transcript level).

### Figure S1

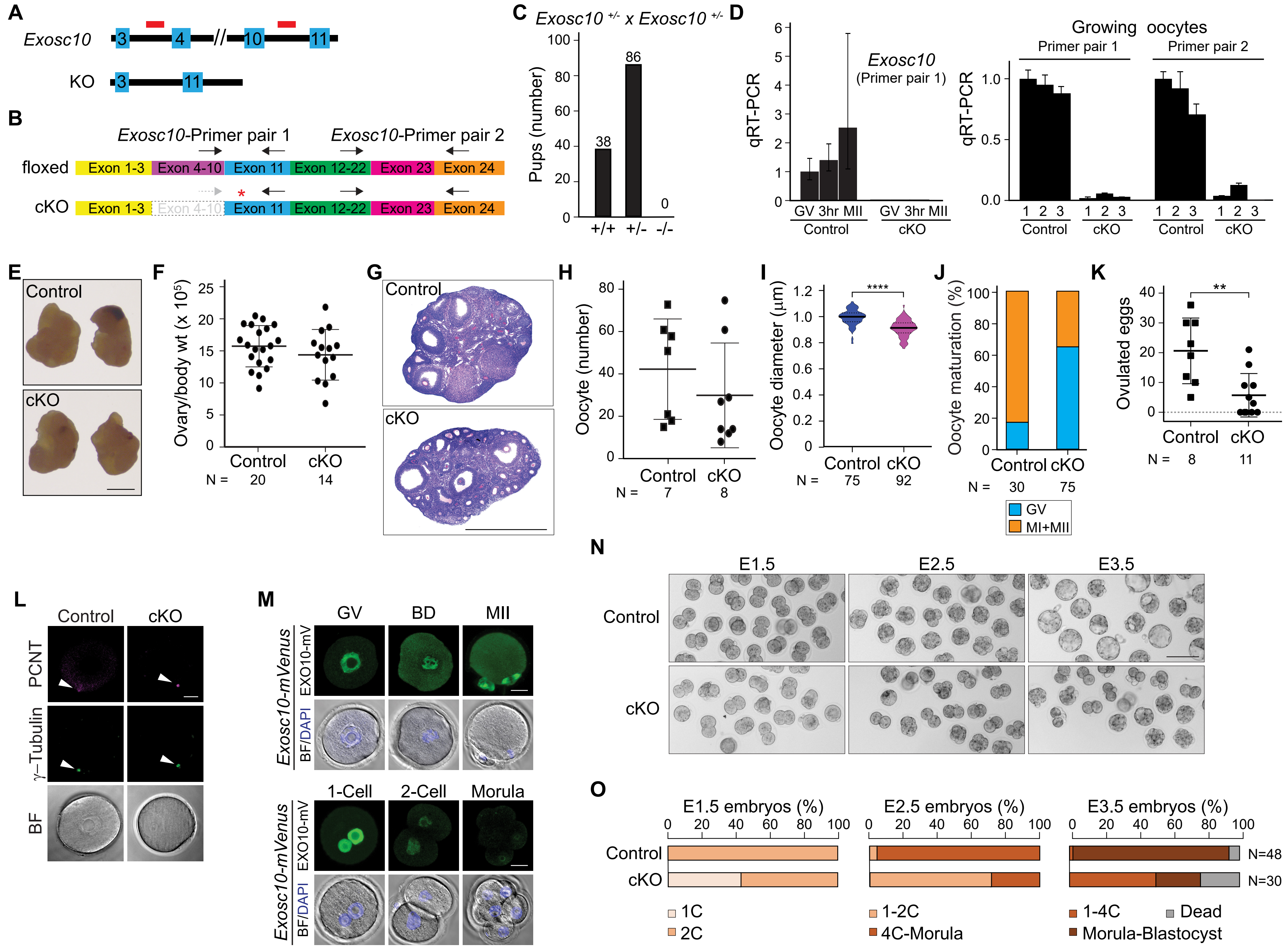

A
Exosc10
3
4
10
11
KO
3
11
B
Exosc10-Primer pair 1
Exosc10-Primer pair 2
floxed
Exon 1-3
Exon 4-10
Exon 11
Exon 12-22
Exon 23
Exon 24
cKO
Exon 1-3
Exon 4-10
Exon 11
Exon 12-22
Exon 23
Exon 24
C
Exosc10 +/- x Exosc10 +/-
Pups (number)
38
86
0
+/+
+/-
-/-
D
qRT-PCR
Exosc10
(Primer pair 1)
GV 3hr MII
GV 3hr MII
Control
cKO
Growing oocytes
Primer pair 1
Primer pair 2
qRT-PCR
1 2 3
1 2 3
1 2 3
1 2 3
Control
cKO
Control
cKO
E
Control
cKO
F
Ovary/body wt (x 10^5)
Control
cKO
N = 20 14
G
Control
cKO
H
Oocyte (number)
Control
cKO
N = 7 8
I
Oocyte diameter (µm)
****
Control cKO
N = 75 92
J
Oocyte maturation (%)
Control cKO
N = 30 75
GV
MI+MII
K
Ovulated eggs
**
Control
cKO
N = 8 11
L
Control
cKO
PCNT
γ–Tubulin
BF
M
GV
BD
MII
Exosc10-mVenus
EXO10-mV
BF/DAPI
1-Cell
2-Cell
Morula
Exosc10-mVenus
EXO10-mV
BF/DAPI
N
E1.5
E2.5
E3.5
Control
cKO
O
E1.5 embryos (%)
E2.5 embryos (%)
E3.5 embryos (%)
Control
cKO
N=48
N=30
1C
2C
1-2C
4C-Morula
1-4C
Morula-Blastocyst
Dead

### Figure S2

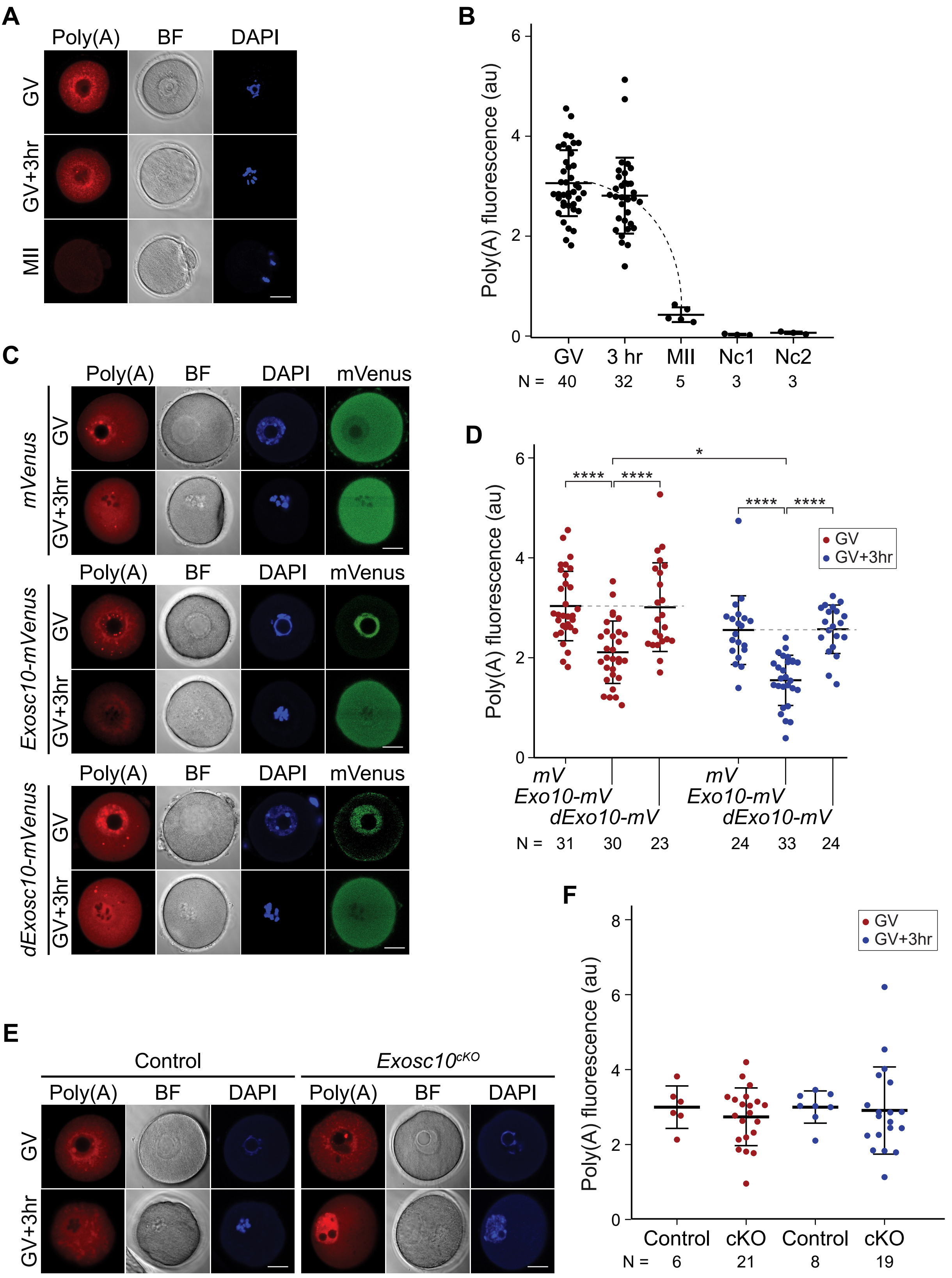
A
Poly(A)
BF
DAPI
GV
GV+3hr
MII
B
Poly(A) fluorescence (au)
GV
3 hr
MII
Nc1
Nc2
N = 40 32 5 3 3
C
Poly(A)
BF
DAPI
mVenus
mVenus
Exosc10-mVenus
dExosc10-mVenus
GV
GV+3hr
D
Poly(A) fluorescence (au)
GV
GV+3hr
mV
Exo10-mV
dExo10-mV
N = 31 30 23 24 33 24
E
Control
Exosc10cKO
Poly(A)
BF
DAPI
GV
GV+3hr
F
Poly(A) fluorescence (au)
GV
GV+3hr
Control
cKO
N = 6 21 8 19

### Figure S3

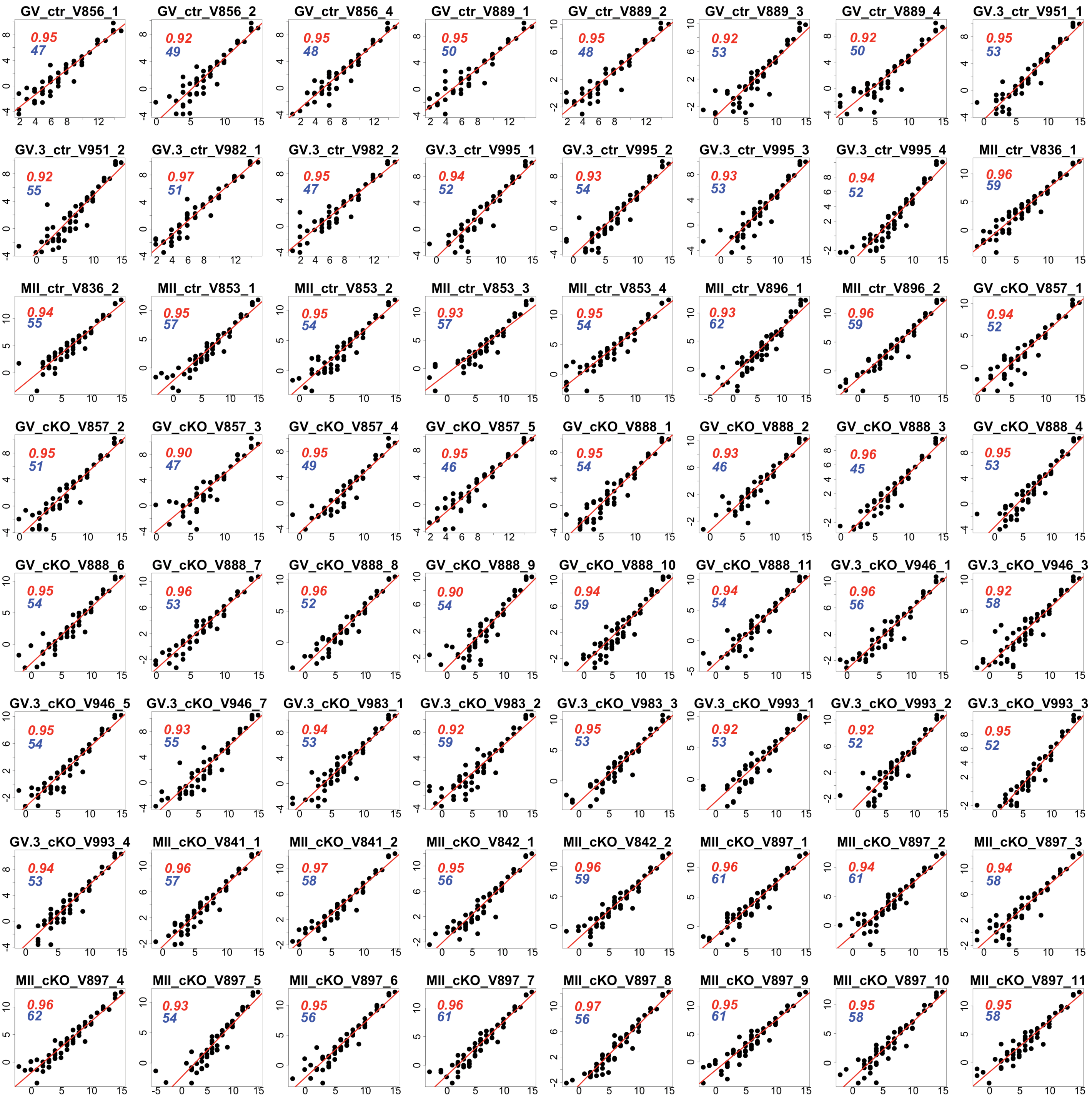
GV_ctr_V856_1 0.95 47
GV_ctr_V856_2 0.92 49
GV_ctr_V856_4 0.95 48
GV_ctr_V889_1 0.95 50
GV_ctr_V889_2 0.95 48
GV_ctr_V889_3 0.92 53
GV_ctr_V889_4 0.92 50
GV.3_ctr_V951_1 0.95 53
GV.3_ctr_V951_2 0.92 55
GV.3_ctr_V982_1 0.97 51
GV.3_ctr_V982_2 0.95 47
GV.3_ctr_V995_1 0.94 52
GV.3_ctr_V995_2 0.93 54
GV.3_ctr_V995_3 0.93 53
GV.3_ctr_V995_4 0.94 52
MII_ctr_V836_1 0.96 59
MII_ctr_V836_2 0.94 55
MII_ctr_V853_1 0.95 57
MII_ctr_V853_2 0.95 54
MII_ctr_V853_3 0.93 57
MII_ctr_V853_4 0.95 54
MII_ctr_V896_1 0.93 62
MII_ctr_V896_2 0.96 59
GV_cKO_V857_1 0.94 52
GV_cKO_V857_2 0.95 51
GV_cKO_V857_3 0.90 47
GV_cKO_V857_4 0.95 49
GV_cKO_V857_5 0.95 46
GV_cKO_V888_1 0.95 54
GV_cKO_V888_2 0.93 46
GV_cKO_V888_3 0.96 45
GV_cKO_V888_4 0.95 53
GV_cKO_V888_6 0.95 54
GV_cKO_V888_7 0.96 53
GV_cKO_V888_8 0.96 52
GV_cKO_V888_9 0.90 54
GV_cKO_V888_10 0.94 59
GV_cKO_V888_11 0.94 54
GV.3_cKO_V946_1 0.96 56
GV.3_cKO_V946_3 0.92 58
GV.3_cKO_V946_5 0.95 54
GV.3_cKO_V946_7 0.93 55
GV.3_cKO_V983_1 0.94 53
GV.3_cKO_V983_2 0.92 59
GV.3_cKO_V983_3 0.95 53
GV.3_cKO_V993_1 0.92 53
GV.3_cKO_V993_2 0.92 52
GV.3_cKO_V993_3 0.95 52
GV.3_cKO_V993_4 0.94 53
MII_cKO_V841_1 0.96 57
MII_cKO_V841_2 0.97 58
MII_cKO_V842_1 0.95 56
MII_cKO_V842_2 0.96 59
MII_cKO_V897_1 0.96 61
MII_cKO_V897_2 0.94 61
MII_cKO_V897_3 0.94 58
MII_cKO_V897_4 0.96 62
MII_cKO_V897_5 0.93 54
MII_cKO_V897_6 0.95 56
MII_cKO_V897_7 0.96 61
MII_cKO_V897_8 0.97 56
MII_cKO_V897_9 0.95 61
MII_cKO_V897_10 0.95 58
MII_cKO_V897_11 0.95 58

### Figure S4

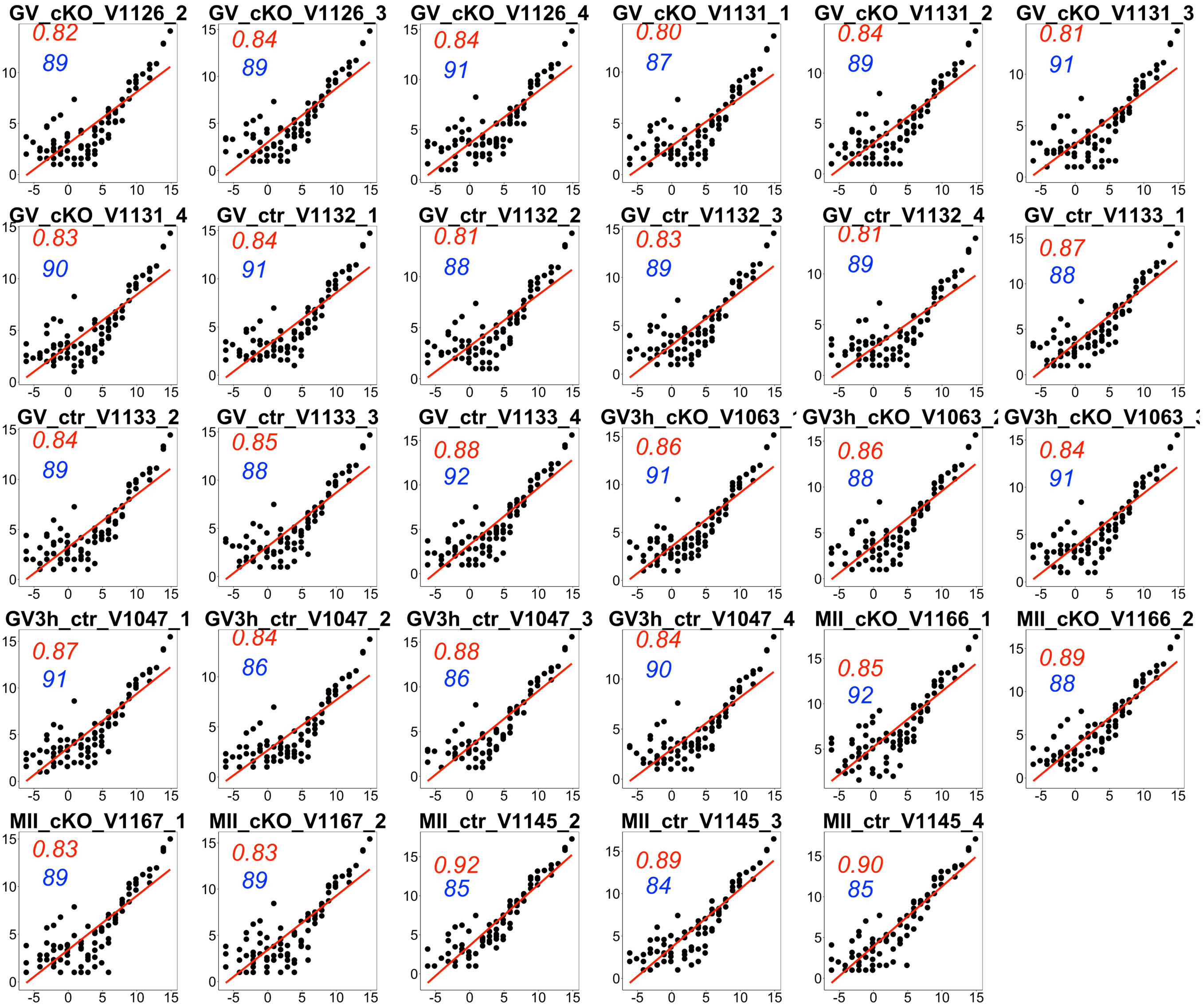
GV_cKO_V1126_2
0.82
89
GV_cKO_V1126_3
0.84
89
GV_cKO_V1126_4
0.84
91
GV_cKO_V1131_1
0.80
87
GV_cKO_V1131_2
0.84
89
GV_cKO_V1131_3
0.81
91
GV_cKO_V1131_4
0.83
90
GV_ctr_V1132_1
0.84
91
GV_ctr_V1132_2
0.81
88
GV_ctr_V1132_3
0.83
89
GV_ctr_V1132_4
0.81
89
GV_ctr_V1133_1
0.87
88
GV_ctr_V1133_2
0.84
89
GV_ctr_V1133_3
0.85
88
GV_ctr_V1133_4
0.88
92
GV3h_cKO_V1063_
0.86
91
GV3h_cKO_V1063_
0.86
88
GV3h_cKO_V1063_
0.84
91
GV3h_ctr_V1047_1
0.87
91
GV3h_ctr_V1047_2
0.84
86
GV3h_ctr_V1047_3
0.88
86
GV3h_ctr_V1047_4
0.84
90
MII_cKO_V1166_1
0.85
92
MII_cKO_V1166_2
0.89
88
MII_cKO_V1167_1
0.83
89
MII_cKO_V1167_2
0.83
89
MII_ctr_V1145_2
0.92
85
MII_ctr_V1145_3
0.89
84
MII_ctr_V1145_4
0.90
85

#### Figure S5

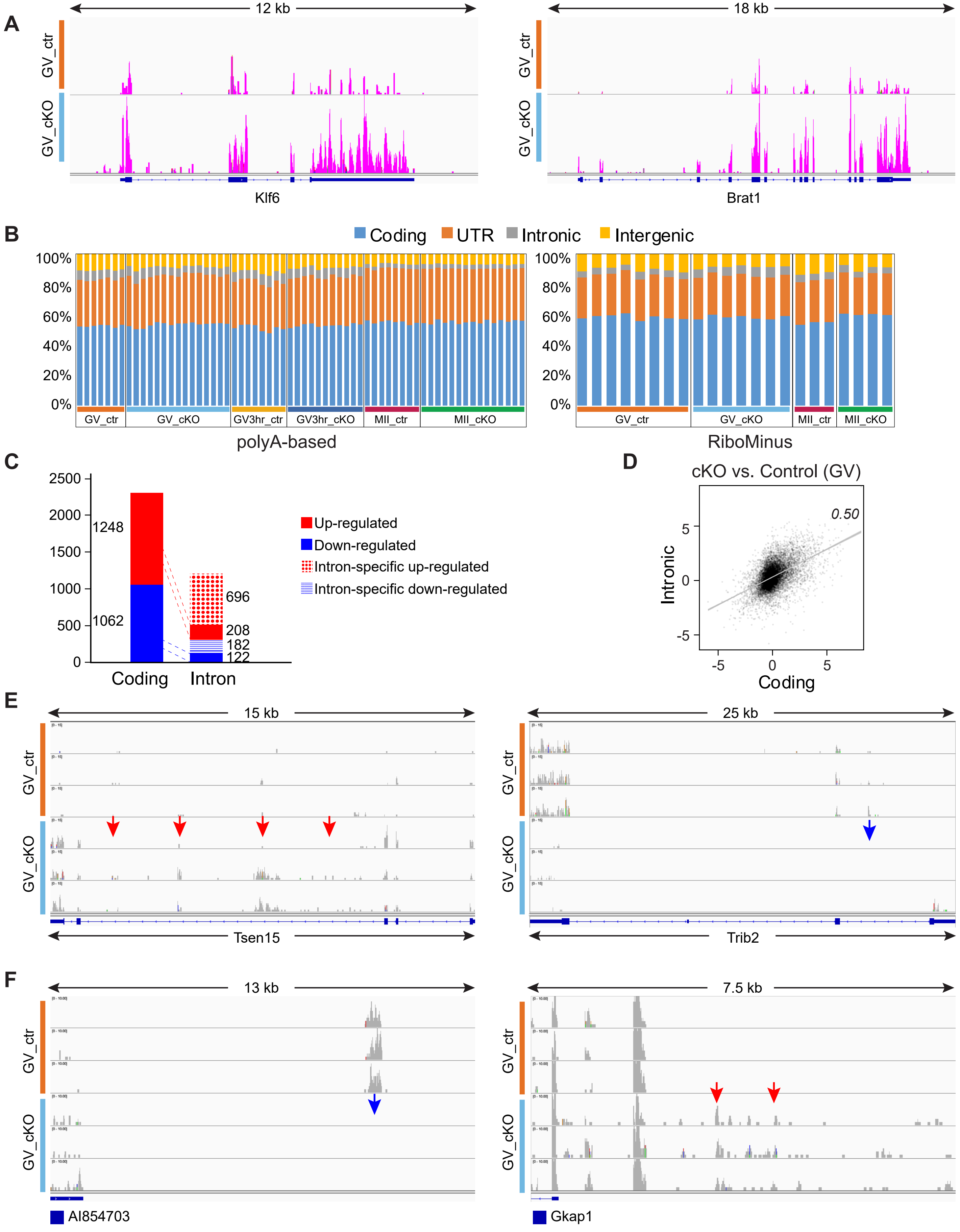

A
12 kb
GV_ctr
GV_cKO
Klf6
18 kb
Brat1
B
Coding
UTR
Intronic
Intergenic
100%
80%
60%
40%
20%
0%
GV_ctr
GV_cKO
GV3hr_ctr
GV3hr_cKO
MII_ctr
MII_cKO
polyA-based
RiboMinus
C
2500
2000
1500
1000
500
0
1248
1062
696
208
182
122
Coding
Intron
Up-regulated
Down-regulated
Intron-specific up-regulated
Intron-specific down-regulated
D
cKO vs. Control (GV)
0.50
Intronic
Coding
5
0
-5
E
15 kb
Tsen15
25 kb
Trib2
F
13 kb
AI854703
7.5 kb
Gkap1

#### Figure S6

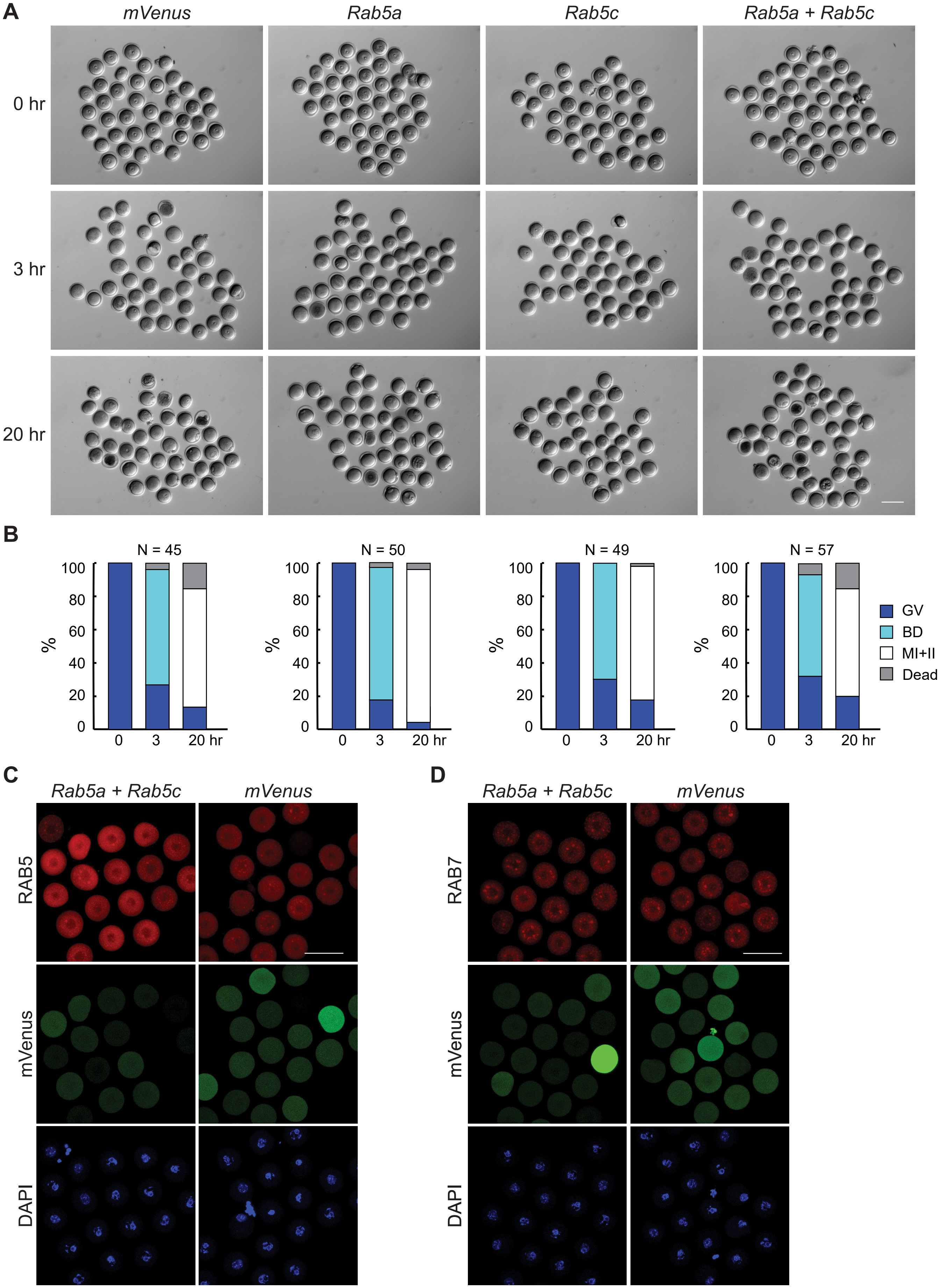
A
mVenus
Rab5a
Rab5c
Rab5a + Rab5c
0 hr
3 hr
20 hr
B
N = 45
N = 50
N = 49
N = 57
100
80
60
40
20
0
%
0
3
20 hr
GV
BD
MI+II
Dead
C
Rab5a + Rab5c
mVenus
RAB5
mVenus
DAPI
D
Rab5a + Rab5c
mVenus
RAB7
mVenus
DAPI

### Figure S7

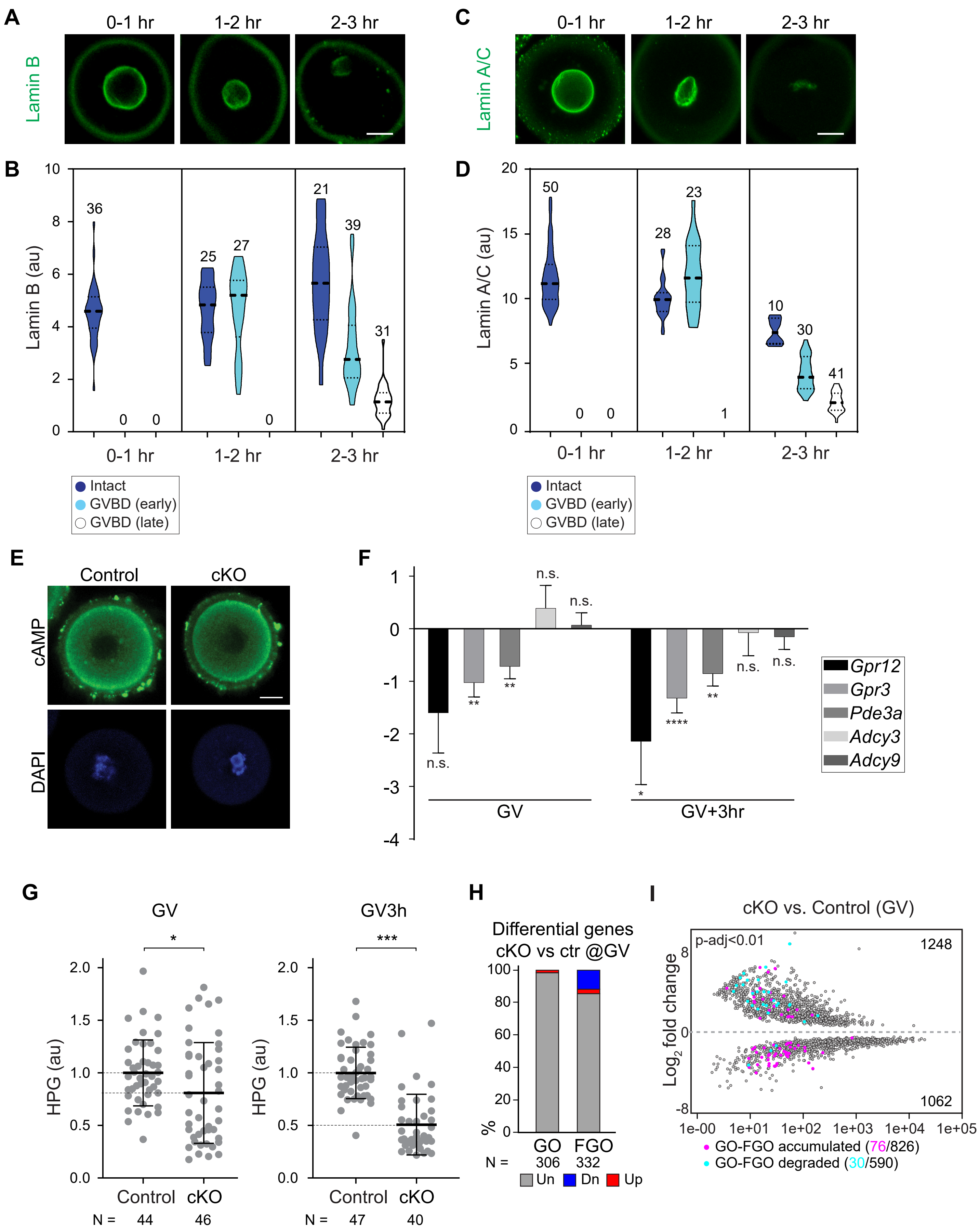

A
0-1 hr
1-2 hr
2-3 hr
Lamin B
C
Lamin A/C
B
Lamin B (au)
36
25
27
21
39
31
0
0
0
Intact
GVBD (early)
GVBD (late)
D
Lamin A/C (au)
50
28
23
10
30
41
1
E
Control
cKO
cAMP
DAPI
F
n.s.
**
****
*
GV
GV+3hr
Gpr12
Gpr3
Pde3a
Adcy3
Adcy9
G
GV
GV3h
HPG (au)
Control
cKO
N = 44 46
N = 47 40
***
H
Differential genes
cKO vs ctr @GV
%
GO
FGO
N = 306 332
Un
Dn
Up
I
cKO vs. Control (GV)
p-adj<0.01
1248
1062
Log2 fold change
GO-FGO accumulated (76/826)
GO-FGO degraded (30/590)
